# Chromatin Remodeling in Patient-Derived Colorectal Cancer Models

**DOI:** 10.1101/2022.07.24.501300

**Authors:** Kun Xiang, Ergang Wang, Gabrielle Rupprecht, John Mantyh, Marcos Negrete, Golshid Sanati, Carolyn Hsu, Peggy Randon, Anders Dohlman, Kai Kretzschmar, Nicholas Giroux, Shengli Ding, Lihua Wang, Jorge Prado Balcazar, Qiang Huang, Pasupathi Sundaramoorthy, Rui Xi, Shannon Jones McCall, Zhaohui Wang, Yubin Kang, Scott Kopetz, Gregory E. Crawford, Hans Clevers, David Hsu, Xiling Shen

## Abstract

Patient-Derived Organoids (PDO) and Xenografts (PDX) are the current gold standards for patient derived models of cancer (PDMC). Nevertheless, how patient tumor cells evolve in these models and the impact on drug response remains unclear. Herein, we compared the transcriptomic and chromatin accessibility landscapes of six matched sets of colorectal cancer (CRC) PDO, PDX, PDO-derived PDX (PDOX), and original patient tumors (PT) and discovered two major remodeling axes. The first axis delineates PDX and PDO from PT, and the second axis distinguishes PDX and PDO. PDOX were more similar to PDX than they were to PDO, indicating that the growth environment is a driving force for chromatin adaptation. Using bivariate genomic footprinting analysis, we identified transcription factors (TF) that differentially bind to open chromatins between matched PDO and PDOX. Among them, KLF14 and EGR2 footprints were enriched in all six PDOX relative to matched PDO, and silencing of KLF14 or EGR2 promoted tumor growth. Furthermore, EPHA4, a shared downstream target gene of KLF14 and EGR2, altered tumor sensitivity to MEK inhibitor treatment. Altogether, patient-derived CRC cells undergo both common and distinct chromatin remodeling in PDO and PDX/PDOX, driven largely by their respective microenvironments, which results in differences in growth and drug sensitivity and needs to be taken into consideration when interpreting their ability to predict clinical outcome.

## Introduction

Patient-derived models of cancer (PDMC) – such as PDX and PDO – are emerging as powerful avatars of patient tumors for pre-clinical therapeutic development (*1-4*). Compared to cell lines and genetically engineered mouse models, PDO and PDX tend to capture more clinical diversity in terms of disease stage and genetic background (*5*). PDX and PDO recapitulate drug sensitivity associated with genetic mutations (*6, 7*) and correlate with clinical response to chemotherapy (*8, 9*). In therapeutic development, PDX and PDO also help researchers address the high failure rates for new cancer drugs in clinical trials (*10, 11*). However, it remains unclear whether tumor cells undergo changes in PDMC compared to the tumor they were derived from and how the distinct microenvironments of these models may impact patient tumor cell growth and drug sensitivity.

CRC is the 3^rd^ most common cause of cancer-related death worldwide with approximately 150,000 new cases diagnosed in the United States annually (*12, 13*). PDX and PDO have been heavily used as pre-clinical models for CRC research and drug development (*14-17*). Recent high-profile studies from multiple groups consistently showed that PDO can predict CRC patient response to chemotherapy and chemoradiation (*18-20*). However, although molecular signatures of the original tumors are largely recapitulated in those models, CRC cells have been shown to lose their dependency on niche factors as they are passaged (*21*), indicating their gradual adaptation to the new environment. Therefore, it is important to understand how patient CRC cells evolve in PDX and PDO respectively compared to the original patient tumor (PT), which can inform which therapeutic axes in PDMC are transferable to clinical outcomes.

In this National Cancer Institute Patient-Derived Model Consortium-sponsored study, we compared the chromatin accessibility landscapes between matched sets of PT, PDO, PDX, and PDOX. All three models exert chromatin alterations when compared to PT cells, representing a PT-PDMC epigenetic axis. Chromatin alterations in CRC cells are more similar between PDOX and PDX than PDO, indicating that the growth environment of the model exerts strong influence on chromatin adaptation in tumor cells. Distinct chromatin alterations and differential TF binding between PDO and PDOX help further delineate the *in vitro* vs. *in vivo* evolution of CRC, which results in differences in tumor growth and drug sensitivity.

## Results

For this study, we derived matched PDO, PDX, and PDOX from CRC specimens and performed Assay for Transposase-Accessible Chromatin using sequencing (ATAC-seq) and RNA-seq (Figure 1A). Nine surgically resected primary and metastatic CRC specimens from consented patients were collected at Duke BioRepository & Precision Pathology Center (BRPC) (Figure 1B, Supplemental Table 1). Eight PDO were generated and three subcutaneous PDX grew within six months of injection. We also developed six PDOX from the PDO (Figure 1A). In total, we successfully established six matched sample sets (Figure 1B). PDO could be maintained throughout serial passages and cryopreservation, and the CRC identity was confirmed by the maintained protein expression of CDX2 and CK20 (Figure 1C, S1A, B). The xenograft models maintained histopathological characteristics similar to the patient tumors they were derived from, and they were histologically confirmed to be CRC by the H&E staining and IHC expression of CDX2+/CK20+/CK7- (Figure 1C, S1C-E).

**Figure 1.**
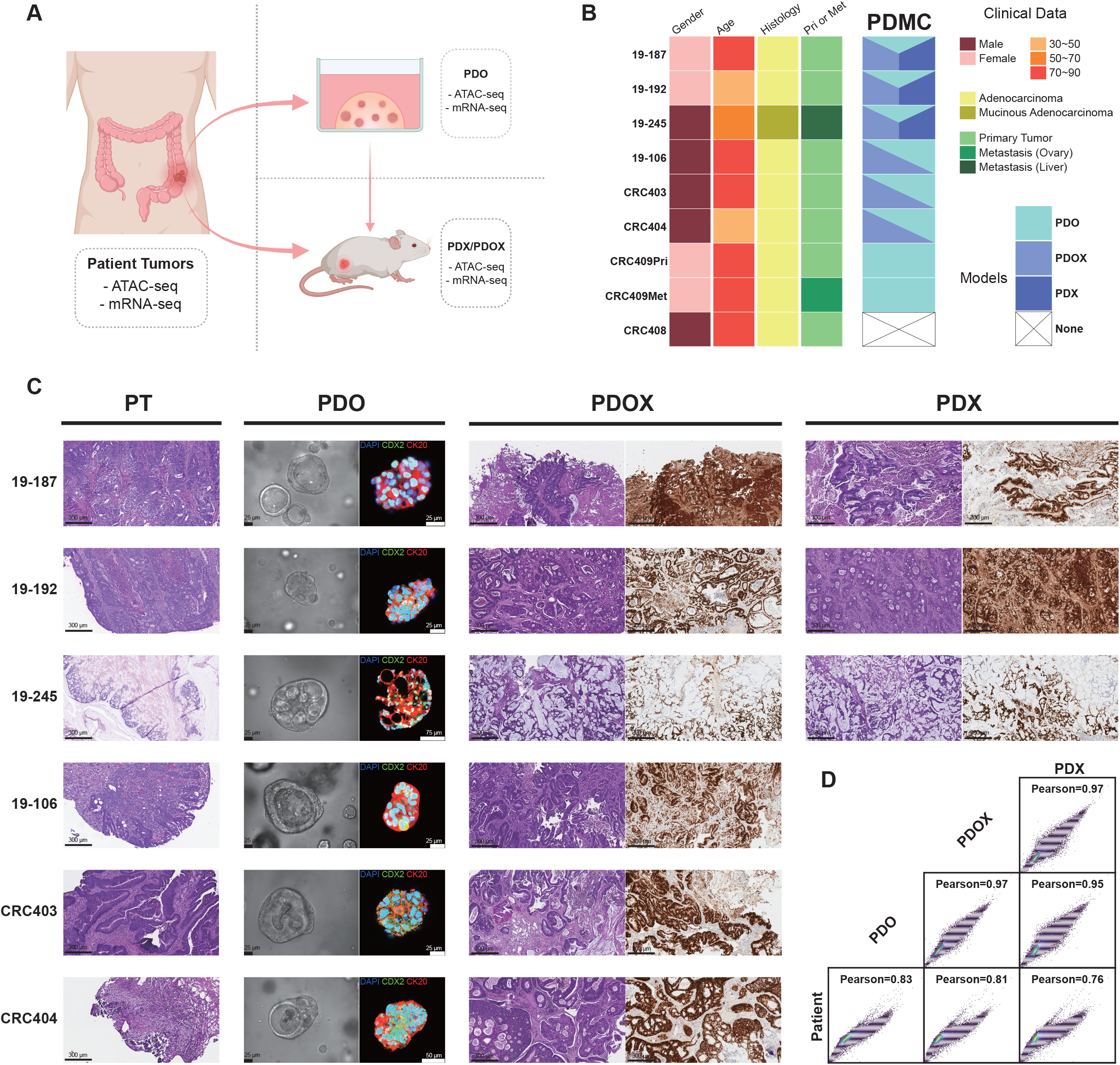
The establishment of matched PT-PDMC sets. **(A)** The overall scheme of the CRC specimens and PDMC processing. The same surgically removed tumor specimens were used to derive both PDO and PDX. PDO were subcutaneously injected into mice to obtain PDOX. All available models, including the original patient tumors, were processed for ATAC-seq and mRNA-seq. **(B)** The summary of model establishment and clinical data for all patients. **(C)** Histology of PT and PDMC, showing the sets of 19-187, 19-192, 19-245, 19-106, CRC403, and CRC404 (the rest are shown in the Supplemental Figure S1). PT panel: H&E-staining of PT at 10X magnification showing the tumor region. Scale bars, 300 µm; PDO panel: The bright-field images (left) of PDO. Scale bars, 25 µm. The confocal images (right) of PDO immunolabeled for CDX2 (green), CK20 (red), and DAPI (blue); PDX panel and PDOX panel: H&E-staining (left) and protein immunostaining for CDX2 (right) of PDX/PDOX. Scale bars, 300 µm. **(D)** Pairwise ATAC-seq correlation between patient samples and models based on normalized read counts of detected peaks. Pearson correlation coefficient is presented for each comparison.

We next profiled the open-chromatin landscapes of the six matched PT-PDMC sets using ATAC-seq (*22, 23*) supplemented by messenger RNA sequencing (mRNA-seq). We obtained high-quality ATAC-seq libraries with high consistency across replicates, as reflected by the QC metrics, including overall sequencing depth, mitochondria fractions, and peak numbers (Supplemental Table 2). Mouse components from the xenografts were removed via the mouse-cell-depletion kit prior to the library preparations, so that the average mapping rates to human genome hg19 were > 90% (Supplemental Table 2).

The global chromatin accessibility landscapes revealed by ATAC-seq were largely congruent across PT and PDMC (PDO, PDX, and PDOX), suggesting that CRC cells mostly retained their identity in PDMC (Figure 1D, Figure S2, S3A). Nevertheless, certain chromatin accessibility alterations could be detected by differential expression analysis based on consensus peaks from PDMC and PT, according to Diffbind (*24*), resulting in separate PT and PDMC clusters in the hierarchical heatmap (Figure 2A).

**Figure 2.**
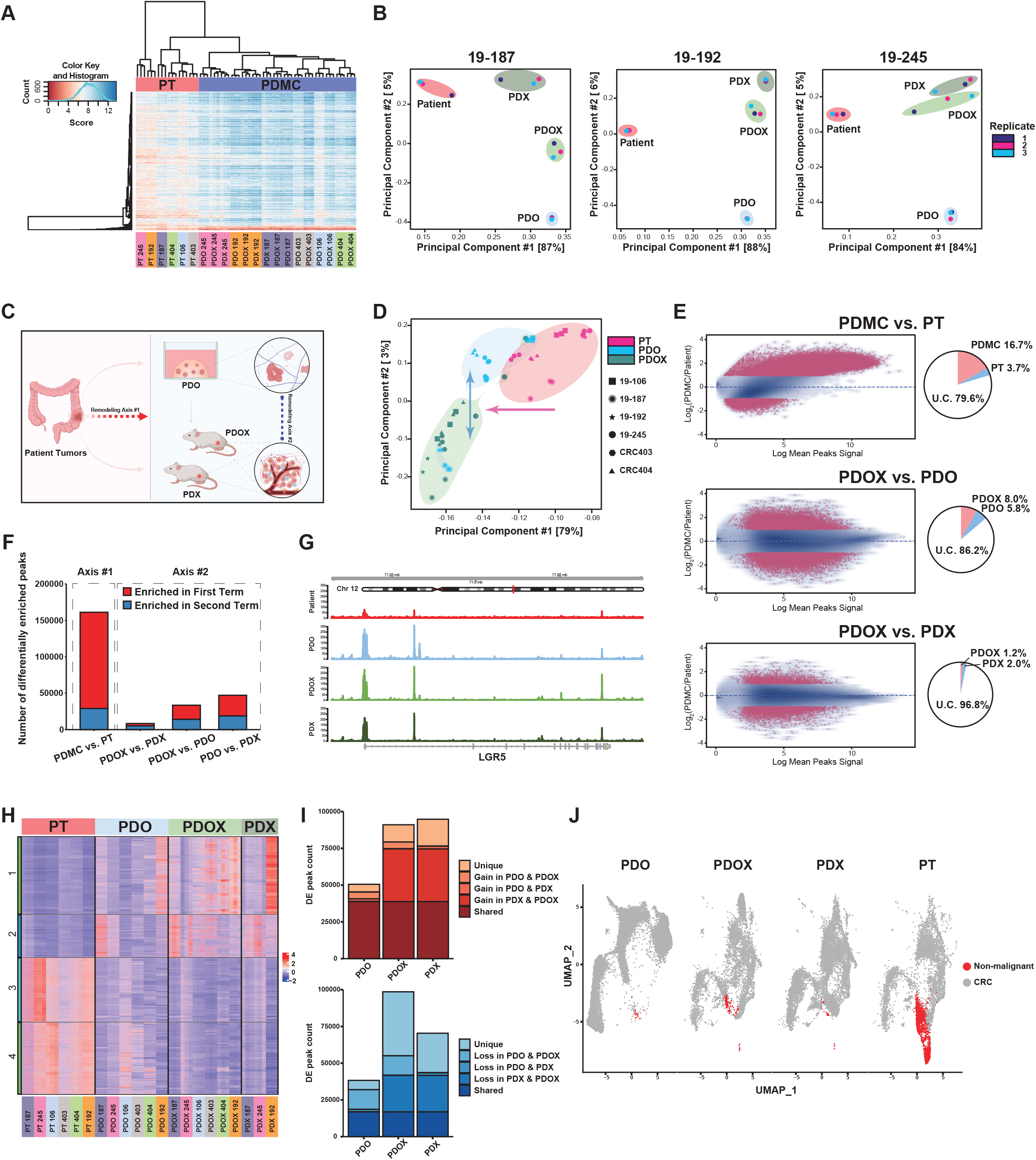
ATAC-seq analysis suggests a two-axes remodeling in CRC cells of PDMC. **(A)** Heatmap of unbiased hierarchical clustering of PT and PDMC based on 1000 chromatin accessible regions with the highest variance. Patient tumor (red) and PDMC (blue) are separated. All replicates are clustered with each other, and the samples from the same patient set are indicated/labeled with a unique color. Six PT-PDMC sets are included: set 245(19-245), set 192 (19-192), set 187 (19-187), set 106 (19-106), set 404 (CRC404), set 403 (CRC403). **(B)** Principal component analysis of individual complete PT-PDMC sets (19-187, 19-192, and 19-245) from ATAC-seq data based on the DiffBind score DBA_SCORE_RPKM. Patient samples are circled in red, PDO in blue, PDX in green, and PDOX in light green. Each type of model has three replicates. **(C)** The illustration of the two-axes remodeling. **(D)** Principal component analysis of the pooled PT-PDO-PDOX sets (*N* = 3 × 6) from ATAC-seq data based on the DiffBind score DBA_SCORE_RPKM. Patient samples are red, PDO are blue, and PDOX are green. Different shapes of dots denote patient IDs. **(E)** Differential analysis of ATAC-seq peaks in the comparisons of PDMC vs. PT, PDOX vs. PDO, and PDOX vs. PDX based on the consensus peak set derived from each condition. Differentially enriched (DE) peaks are labeled in red, |log2(FC)|>1 and p<0.05. Pie charts show the percentages of enriched or unchanged (U.C.) peaks. **(F)** The number of differentially enriched peaks in different model comparisons. The number of peaks enriched in the first or second term of the comparison is denoted (Supplemental Table 3). The remodeling axis #1 includes PDMC vs. PT. The remodeling axis #2 includes PDOX vs. PDX, PDOX vs. PDO, and PDO vs. PDX. **(G)** ATAC-seq signal track showing LGR5 locus in different models. The exon locations are indicated in the gene map. **(H)** Heatmap of all ATAC differentially enriched peaks gained or lost between patient samples, PDO, PDOX and PDX based on the consensus peak set. Each column represents a replicate of ATAC sequencing of PT or models. **(I)** The number of ATAC-seq peaks that are significantly gained (top) or lost (bottom) in PDMC versus PT samples, shared or unique to models. **(J)** Anchored single cell multiome analysis for PDMC and PT, visualizations of CRC and non-malignant components in the PDMC and PT samples.

Principal component analysis (PCA) of the chromatin accessibility showed clear separation between PT, PDO, and PDX/PDOX (Figure 2B, S4A). In all six cases, the first principal component (PC) axis distinguished PT from all PDMC, and the second PC axis distinguished *in vivo* models (PDX/PDOX) from the *in vitro* model (PDO) (Figure 2C). PDOX were more similar to PDX than PDO along the second axis. The pooled six PT-PDMC sets revealed that the two epigenetic axes were still largely preserved despite patient-to-patient variation (Figure 2D), which was also reflected in transcriptomic PCA from mRNA-seq data (Figure S4B). PDOX and PDO were also highly correlated versus PT according to both ATAC-seq and RNA-seq (Figure S4C, D), in concordance with the first PCA axis.

Although the majority of consensus peaks remained unchanged (79.6%), paired differential analysis identified alterations in chromatin accessibility between PT and PDMC (Figure 2E, F, S5A, B), such as at loci in chromosomes 7, 8, 13, and X (Figure S3B). For instance, there were considerable alterations between PDMC and PT at the locus associated with LGR5, a stem cell and proliferation marker for colon stem cells and CRC (*25-28*) (Figure 2G). Compared to PT vs. PDMC along the first axis, fewer differentially enriched peaks were detected for PDOX vs. PDO and PDOX vs. PDX along the second axis (Figure 2E, F, S5A, B). The differentially enriched peaks were fewest between PDX and PDOX (Figure 2F), in concordance with the PCA analyses (Figure 2B). PDO, PDOX, and PDX had similar overall chromatin accessibilities at the chromosome level, but there were more differences for PDOX vs. PDO than PDOX vs. PDX.

We performed differential chromatin accessibility analysis on the consensus ATAC-seq peak set across the matched sets of PT, PDO, PDOX, and PDX. Unsupervised hierarchical clustering of all differentially accessible peaks identified gain/loss clusters that are shared or unique among PDMC (Figure 2H, I). KEGG pathway and Gene Ontology analyses based on gain/loss peaks further suggest pathways that may be commonly or uniquely altered among PDMC (Figure S6A-C). The pathway analysis on the shared loss peaks suggests PT likely contains more stromal cells. We performed 10X Single Cell Multiome (RNA+ATAC) sequencing on one set of PT-PDO-PDOX-PDX (Figure S7A). The stromal component is higher in PT than in PDMC (Figure 2J, S7B), which may have contributed to the difference between PT and PDMC.

The findings that PDOX were more similar to PDX than PDO (Figure 2F, H, I) suggests that chromatin alterations were largely driven by the tumor environment. Therefore, direct comparison of PDO and PDOX may reveal how the *in vivo* environment remodels the chromatin of CRC cells from PDO. KEGG pathways analysis (Figure S8) suggested pathways involving interactions with extracellular matrix and signaling transductions (BRAF, MAPK, EPH-Ephrin) were altered between PDO and PDOX. To investigate potential transcription factors (TFs) that may regulate this process, we performed bivariate genomic footprinting (BaGFoot) analysis, which is based on the changes in the depth of TF footprint and TF motif-flanking accessibility (*29*). 57 TFs presented higher activities in PDOX, and 47 TFs exhibited higher activities in PDO (Figure 3A, Supplemental Table 4). Several of these TFs were also identified when comparing PDO vs. PT and PDOX vs. PT, respectively (Figure S9A), including EVX2, reported to be methylated in lung cancer (*30*), and SNAI1/2, involved in epithelial-to-mesenchymal transitions (EMT) and responsive to EGF (*31, 32*). These TF footprints were further analyzed by TOBIAS BINDetect (*33*) (Figure S9B),

**Figure 3.**
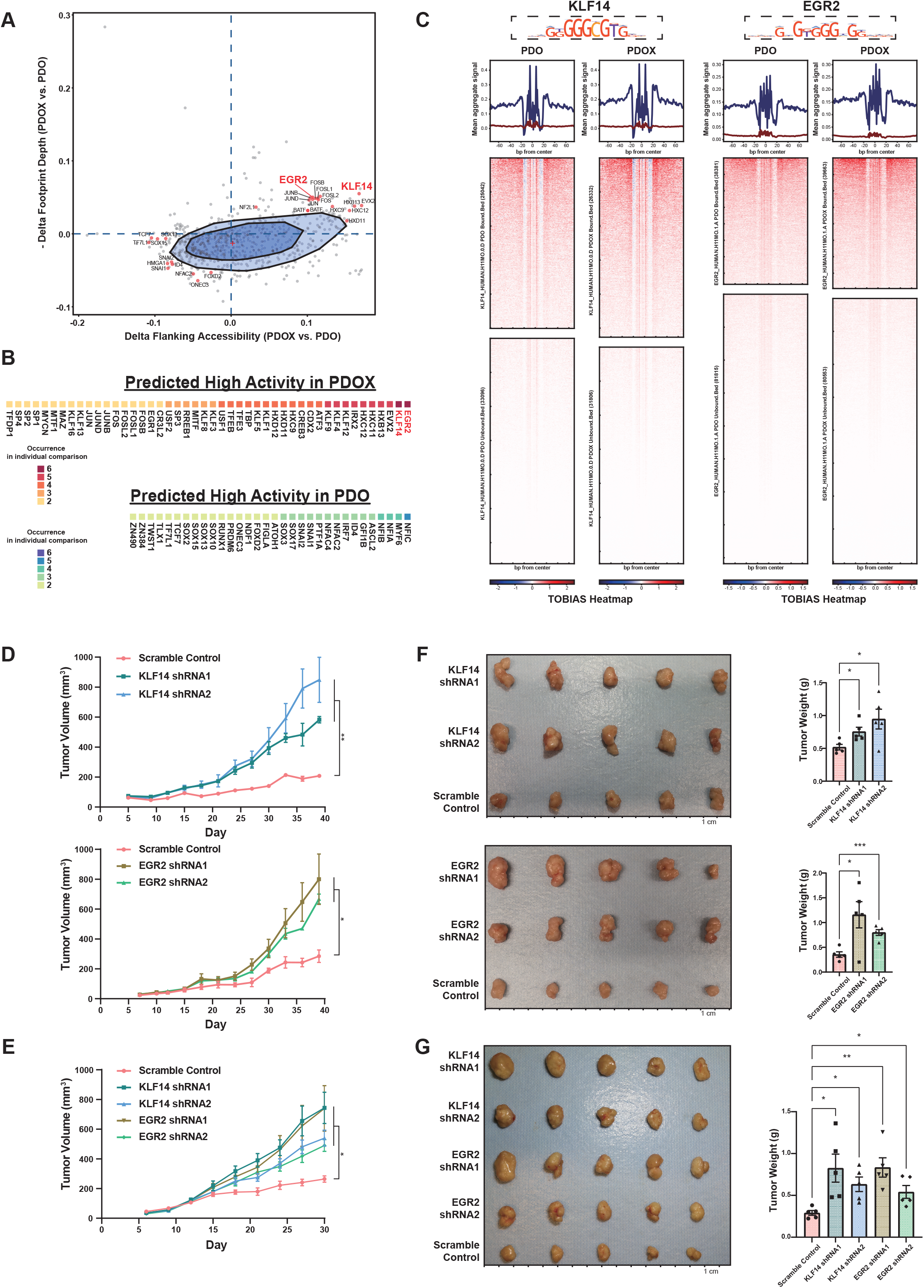
Transcription factor activities are affected by PDO->PDOX remodeling. **(A)** BaGFoot analysis of PDOX vs. PDO based on consensus peak set derived from each condition (*N* = 3 × 6). The TFs predicted to be active in PDOX are in the first quadrant beyond the fence area with an extending factor of 1.5. The model specific TFs (also predicted to be active in PDOX in the comparison of PDOX vs. PT, Supplemental Table 4), as well as EGR2 and KLF14, are highlighted. The TFs in the third quadrant beyond the fence area are predicted to be active in PDO. The model specific TFs (also predicted active in PDO in the comparison of PDO vs. PT, Supplemental Table 4) are highlighted. **(B)** The occurrence counts of high activity (beyond the fence in the first or third quadrant) TFs in PDOX vs. PDO from BaGFoot analysis for individual patient sets. **(C)** TOBIAS footprint analysis of KLF14 (motif: KLF14_HUMAN.H11MO.0.D) and EGR2 (motif: EGR2_HUMAN.H11MO.1.A). The aggregated signals are summarized at the top; blue lines are from bound sites, and red lines are from unbound sites. The footprint heatmaps indicate the chromatin accessibility in the bound and unbound sites of KLF14 and EGR2 in PDO and PDOX. **(D-E):** Tumor growth curves of PDOX 19-106 (**D**) and PDOX 19-187 (**E**) for KLF14 and EGR2 shRNA KD or the scrambled control. Error bars denote SEM of five replicates. p values were calculated based on repeat measurement one-way ANOVA with Fisher’s LSD test. *p<0.05, **p<0.01. (**F-G**): Photos of xenograft tumors (left) and tumor weight comparisons (right) of PDOX 19-106 (**F**) and PDOX 19-187 (**G**) for KLF14 and EGR2 KD and the scramble control. The ruler of the tumor photo has 1-cm intervals. Error bars in the tumor weight plots denote SEM of five replicates (mouse replicates are shown as scatter dots). p values were calculated based on ANOVA with post-hoc. *p<0.05, **p < 0.01, ***p < 0.001.

We then performed BaGFoot analysis for each individual patient set (Figure S10A, Supplemental Table 4). Notably, two TFs, Krüppel-like family 14 (KLF14) and Early Growth Response 2 (EGR2), stood out in all six patient sets (Figure 3B, S10B). TOBIAS footprint analysis confirmed that KLF14 and EGR2 had deeper footprints in PDOX than PDO (Figure 3C, S10C). Krüppel-like family factors have been reported to regulate a multitude of cancer-relevant processes (*34*). KLF14 precludes KRAS-associated cell growth and transformation (*34, 35*), and loss of KLF14 can trigger PIK4-mediated centrosome amplification to promote colon tumorigenesis (*36*). KLF14 has also been associated with high-density lipoprotein cholesterol levels and metabolic syndrome (*37*). EGR2 transactivates BNIP3L/BAK in PTEN-induced apoptosis (*38*) and binds ARF promoters and p16 to induce senescence (*39*). EGR2 malfunctions are also known to be related to cancer (*40, 41*).

To assess their functions, we knocked down KLF14 and EGR2 in both PDO 19-106 and 19-187 using lentiviral shRNA and injected the PDO into immune-deficient NSG mice to form PDOX (Figure S11A). For each gene, we used two shRNAs along with a scrambled shRNA as the control (Figure S11B). KLF14- and EGR2-deficient PDOX grew faster than PDOX with scramble control shRNA (Figure 3D, E). We collected all xenograft tumors at the end of the experiments. For 19-106, the final tumor masses were 1.46- and 1.83-fold higher in KLF14-deficient PDOX and 3.26- and 2.25-fold higher in EGR2-deficient PDOX compared to the scrambled control group (Figure 3F). For 19-187, the final tumor masses of KLF14-shRNA1, KLF14-shRNA2, EGR2-shRNA1, and EGR2-shRNA2 were 2.84-, 2.18-, 2.87- and 1.90-fold higher than control PDOX, respectively (Figure 3G). Therefore, KLF14 and EGR2 binding activities were upregulated by the *in vivo* environment, and reducing KLF14 and EGR2 resulted in enhanced tumor growth.

We then analyzed the genes downstream of KLF14 and EGR2 by integrating ATAC-seq detected peaks and gene expression (Figure 4A), and identified 24 genes associated with both EGR2 and KLF14 that are upregulated in PDOX (Supplemental Table 5). Among them, KLF4 and EPH Receptor A4 (EPHA4) are the top two candidates. Since KLF4 is a TF just like KLF14 and EGR2, we focused on EPHA4, whose KLF14 and EGR2 binding regions are more accessible in PDOX than in PDO in all 6 cases, consistent with higher EPHA4 gene expression in PDOX (Figure 4B-D, S12A). Additionally, single cell multiomics data link noncoding accessible DNA elements to the expression of the gene (*42*). The elevated expression of EPHA4 in PDOX putatively results from the collective regulation of the promoter and three other enhancers, where the promoter has the highest correlation score (Figure 4E). EPHA4 has been reported to interact with fibroblast growth factor receptors (FGFRs) and influences MAPK and AKT pathways (*43-45*). To examine whether KLF14 and EGR2 mediated chromatin remodeling of EPHA4 affects drug sensitivity, we knocked down EPHA4 in the PDO (Figure S12B) then evaluated their responses to 5-Fluorouracil (5FU, chemotherapy), Pemigatinib, Erdafitinib (FGFR inhibitors), and Mirdametinib (MEK inhibitor). Silencing of EPHA4 did not change PDO sensitivity to 5FU and had minor effects with regard to FGFR inhibitors; however, silencing of EPHA4 sensitized PDO to Mirdametinib significantly (Figure 4F, S12C). In addition, we performed a screen of 147 FDA-approved anticancer compounds. Altered EPHA4 expressions influenced PDO responses to some of the drugs (Figure 4G). Therefore, the differential binding of KLF14 and EGR2 and expression of downstream EPHA4 impact both tumor growth and drug sensitivity, suggesting that chromatin remodeling in different PDMC may interfere with their ability to predict therapeutic outcomes.

**Figure 4.**
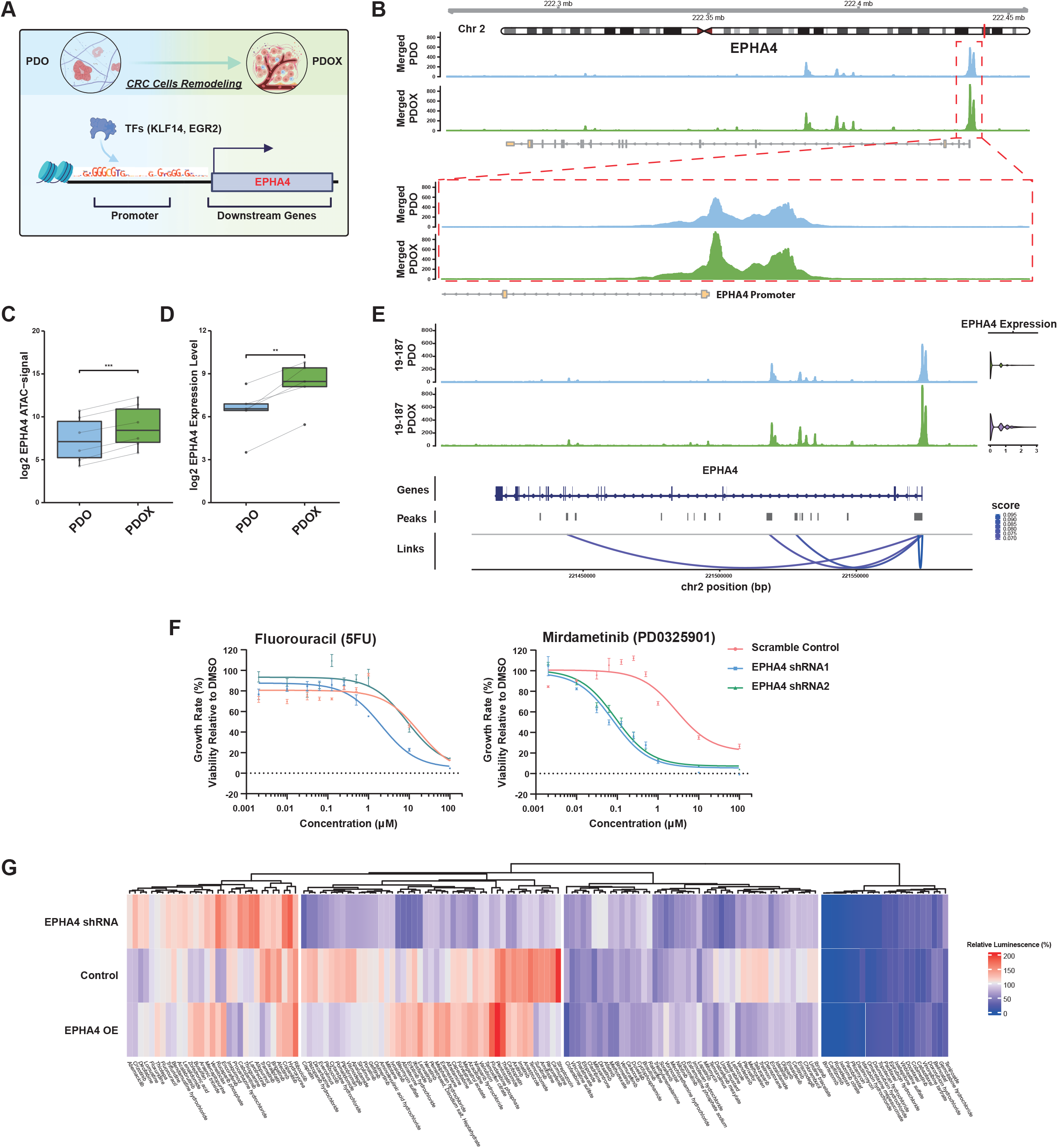
Downstream gene EPHA4 affects drug sensitivities. **(A**) The illustration shows the epigenetic reprogramming of CRC cells from PDO in vitro to PDOX in vivo, where the enhanced binding activities of TFs (KLF14 and EGR2) would lead to the higher expression of the downstream genes (EPHA4). **(B)** ATAC-seq signal track showing EPHA4 locus in PDO and PDOX (all six patient sets merged). The exon locations are indicated in the gene map. The promoter areas of EPHA4 are circled and presented in the bottom panel. **(C)** The boxplot reports cumulative ATAC-seq signals in the EPHA4 region in all paired patient sets of PDOX vs. PDO, p values were calculated based on paired Student’s t-test. ***p<0.001. **(D)** The boxplot reports the EPHA4 expression levels based on RNA-seq in paired PDOX vs. PDO. p values were calculated based on paired Student’s t-test. **p<0.01. **(E)** Peak–gene links for EPHA4 based on PDO and PDOX 19-187 single cell multiome data, showing the correlations between DNA accessibilities and EPHA4 expression. **(F)** PDO19-106 growth rate dose response curves to Fluorouracil and Mirdametinib after knocking down EPHA4. Error bars denote SEM of four replicates. **(G)** Heatmap of drug screen on PDO19-106 (control PDO, EPHA4 shRNA PDO, and EPHA4 overexpression [OE] PDO), showing the average relative luminescence (%) of three replicates for each compound and condition.

## Discussion

PDMC are becoming the gold standard pre-clinical models for therapeutic development and precision oncology. However, how tumor cells evolve in PDMC remain largely unclear. In this study, we established six PT-PDO-PDX/PDOX sets from nine CRC specimens. ATAC-seq, supplemented by RNA-seq, provided a comprehensive picture of the changing chromatin accessibility landscape in PDMC from original PT. Notably, all PDMC are separated from PT along the first PC axis, whereas *in vitro* and *in vivo* PDMC are separated along the second axis. The observation that PDOX is more similar to PDX than to PDO indicate that the PDMC environment is a driving force of chromatin remodeling in tumor cells.

As part of the NCI patient-derived model consortium, we and other researchers have consistently observed that PDOX are generally faster to develop, with higher success rates, than PDX. The difference is more significant for cancer types for which the success rates of conventional PDX are low (*46, 47*). However, we do not have a complete understanding of the tradeoffs. The fact that PDOX is similar to PDX in terms of chromatin accessibility suggests that PDOX may provide a reasonable alternative to PDX for certain applications.

A comparison of PDO and PDOX further highlighted the regulatory elements that respond differently to *in vitro* vs. *in vivo* environments. BaGFoot and TOBIAS Footprinting analyses further revealed TFs that differentially bind to open chromatins in the two models. Among these TFs, KLF14 and EGR2 have enriched motif-binding footprints in PDOX from all six patient cases. Silencing of KLF14 and EGR2 led to enhanced CRC growth in PDOX. This suggests that the mouse stroma can elicit tumor suppressors that slow down the tumor growth, which may partially explain why PDX tend to grow slower than PDO. In addition, the differential expressions of downstream genes like EPHA4 may alter sensitivity and resistance to drugs such as targeted therapy. Therefore, chromatin remodeling of different PDMC may interfere with their ability to predict therapeutic outcomes and need to be more carefully examined. This work provides a resource for the PDMC community to examine the differences among CRC PDMC models, and similar studies in other caner types will likely to be informative.

## Funding

This work was supported by NIH U01CA217514 (XS, DH) as part of the NCI Patient-Derived Models of Cancer Consortium. The BioRepository & Precision Pathology Center at Duke (the source of the patient samples) receives support from NIH P30CA014236 and UM1CA239755.

## Author contributions

X. S. and D. H. initiated the idea and project. K. X., E.W., D. H., and X. S. designed the experiments. S. K., and G. C. advised on experimental designs and data analysis. E. W., K. X., A. D., and N. G. analyzed the data. K. X. performed the experiments with the assistance of M. N., G. S., E. W., and C. H.. D. H., G. R., and S. J. M. facilitated patient tumor samples and medical data acquisition. D. H., G. R., P. S., and Y. K. developed PDX models. K.K. and H.C provided supports on the development of PDO. K. X., J. M., and M. N. established PDO and PDOX models and performed related experiments with the assistance of G. S., Z. W., Q. H., C. H., and P. R.. K. X. prepared sequencing libraries with the assistance of G. S., L. W., R. X., and J. B.. S. D. coordinated the Illumina sequencing. K. X., E. W., and X. S. wrote the manuscript.

## Competing interests

The authors declare no competing interests.

## Data and Materials Availability

Supplemental Information includes Materials and Methods, supplemental figures, and supplemental tables, which can be found with this article online. The biological materials are available from the author by request, the sample transfer needs to be complied with the Duke BRPC Policy. The sequencing data can be obtained from Gene Expression Omnibus (GEO: GSE184482).

## Code Availability

No custom code or software was used in the data analysis. All packages used are listed in the Materials and Methods.

## SUPPLEMENTARY MATERIALS

### Materials and Methods

#### Patient Samples

All CRC specimens were collected at Duke University Hospital through the Duke BioRepository & Precision Pathology Center (BRPC) which is part of the National Cancer Institute’s Cooperative Human Tissue Network. The study was reviewed and approved by the Duke Institutional Review Board (Pro000089222). Informed consent was obtained from all participants. The clinical data of patients were collected through the BRPC medical record system. Each surgically removed tumor specimen was minced then split for three purposes: 1) sequencing as the PT samples, 2) deriving the matched PDO (described in the following method), 3) deriving the matched PDX (described in the following method).

#### Tumor Isolation and Patient-derived Organoid Culture

The PDO development and cultivation methods were modified based on the previously published method (*1, 21, 48*). Briefly, the CRC specimen was minced and incubated in digestion buffer (HBSS with 1 mg/mL collagenase from Clostridium Histolyticum, 0.1 mg/mL DNase I, 3 mM calcium chloride, 100 µg/mL primocin, and 10 µM Y-27632) at 37 °C for 60 minutes in the orbital shaker. Large fragments in the mixture were removed by passing through a 100-µm cell strainer after the incubation. The cells were centrifuged and washed twice with PBS. Cells were counted, embedded in ice-cold Matrigel, and inoculated in 24-well plates. After at least 15 min at 37 °C, Matrigel was polymerized. The CRC culture medium (Advanced DMEM/F12 with 2 mM GlutaMAX, 10 mM HEPES, 100 U/mL Penicillin and Streptomycin, 1X B27, 1.25 mM n-Acetyl Cysteine, 10 nM Gastrin I, 500 nM A-83-01, 3 µM SB202190, 100 ng/mL Recombinant Human R-Spondin, 100 ng/mL Recombinant Human Noggin, 50 ng/mL Recombinant Human EGF, 10 nM Prostaglandin E2, 100 µg/mL Primocin, and 10 mM Nicotinamide) (*1, 21*), was added and refreshed every two to three days.

For the PDO that successfully grew and derived, we passed the organoids every 1-2 weeks based on their growth rates. To passage the organoids, we removed the CRC culture medium and degrade the Matrigel dome with PBS. The mixture of PBS with Matrigel-organoids was collected and centrifuged at 1500 RPM for 5 min, and the PBS was removed. TrypLE™ express enzyme (2 – 5 mL, Gibco) was added, and organoids were incubated at 37 °C for 10 minutes. After the addition of PBS, the cells were centrifuged, rewashed with PBS, and seeded with Matrigel in a 24-well plate. Culture medium was added after Matrigel polymerization. The bright-field images were taken by a Leica DMIL LED Fluorescent Microscope. Part of the established PDO was dissociated then cryopreserved for sequencing using frozen medium (70% FBS + 20% culture medium + 10% DMSO) in liquid nitrogen.

#### PDO H&E-staining and Immunolabeling Fluorescence Staining

PDO were fixed and embedded in paraffin per a modified Trevigen, Inc. protocol (*49*). First, Matrigel domes with PDO were washed with PBS and then fixed with 5 mL of 2% paraformaldehyde (PFA) + 0.1% glutaraldehyde (GA) in PBS at room temperature for 30 minutes. After washing with PBS, the domes were taken to 20% sucrose and left overnight at 4 °C until the domes sank to the bottom. Next, the solution was changed to 70% ethanol, and the domes with PDO were embedded with paraffin for sectioning and H&E-staining in Duke Pathology Research Histology Lab.

For the PDO immunolabeling staining, we followed the PDO 3D imaging protocol published previously (*50*). Briefly, the organoids were recovered from Matrigel using the ice-cold cell recovery solution (Corning) and then fixed with 4% PFA at 4 °C for 45 min. After blocking, PDO were cultured with primary and then secondary antibodies for immunolabeling. Fructose–glycerol clearing solution was used for imaging the organoids under the Leica SP5 inverted confocal microscope. The following antibodies were used for immune-fluorescence staining: CDX2 (12306S, Cell Signaling Technology), 1:200; CK20 (60183-1-IG, Proteintech), 1:100; Anti-Rb IgG-488 (ab150061, Abcam), 1:500; Anti-Ms IgG-594 (ab150112, Abcam).

#### Xenograft Development

All animal experiments were approved by the Duke Animal Care and Use Program (IACUC) following the A235-18-10 and A112-18-05 protocols. PDX were developed as described previously (*51, 52*). The CRC tissue was minced and resuspended in 100 µL of PBS, and injected subcutaneously (s.c.) into the flanks of NOD-SCID IL2Rgamma^null^ (NSG, JAX005557) mice. When the tumor reached a volume of 1000 mm^3^, it was harvested, minced, and resuspended in PBS before passaging, s.c. injection into the flanks of other NSG mice. Part of the tumor tissue from PDX was cryopreserved in 70% FBS + 20% DMEM + 10% DMSO for future passages. We also developed organoid xenografts (PDOX) using a modified version of the published protocol (*53*). PDO were harvested by removing Matrigel with TrypLE™ express enzyme, counted, and resuspended in 50% Matrigel/PBS solution. Cells (1 × 10^6^) were injected s.c. into the flanks of NSG mice. The harvested xenograft tumors were dissociated into single cells using the Tumor Dissociation Kit (Miltenyi Biotech, # 130-095-929), and then they were cryopreserved before sequencing using frozen medium (70% FBS + 20% culture medium + 10% DMSO) in liquid nitrogen. For xenograft staining, tumors were harvested, fixed in 10% neutral buffered formalin, and then embedded in paraffin. Sections were subjected to H&E as well as immunohistochemical staining in Duke Pathology Research Histology Lab.

#### shRNA Lentivirus Transduction of PDO

The KLF14, EGR2, and EPHA4 lentivirus constructs were obtained from Sigma Mission shRNA (KLF14 shRNA1: SHCLNG-NM_138693, TRCN0000107420; KLF14 shRNA2: SHCLNG-NM_138693, TRCN0000107424; EGR2 shRNA1: SHCLNG-NM_000399, TRCN0000013840; EGR2 shRNA2: SHCLNG-NM_000399, TRCN0000013841; EPHA4 shRNA1: SHCLNG-NM_004438, TRCN0000010165; EPHA4, shRNA2: SHCLNG-NM_004438, TRCN0000332976; Scramble Control: pLKO scramble shRNA puro, Addgene #1864). The lentiviral vectors were co-transfected with packaging and envelope plasmids (pCMVR8.74, Addgene #22036; pCMV-VSV-G, Addgene #8454) into 293T cells. The viral supernatant was collected and concentrated using Lenti-X Concentrator (TaKaRa, 631232) 48 hours after transfection. The transduction process is from a modified published protocol (*54*). Briefly, PDO were harvested and resuspended in 500 µL CRC culture medium. Next, the organoids solution was mixed with 500 µL viral particles and 5 µL TransDux (system bioscience, LV850A-1). The mixture was transferred to a 12-well plate and centrifuged at ∼30 °C at 600 g for 60 min. Then, the organoids-virus mixture was incubated for another 3 hours at 37 °C before Matrigel embedding and cultured in the CRC culture medium. On day 3 after transduction, puromycin (5 µg/mL for PDO19-106 and 2 µg/mL for PDO19-187) was added to the medium to select the cells carrying constructs of interest. After 2 weeks of selection, the knockdown efficiencies of KLF14, EGR2, and EPHA4 were evaluated by TagMan qPCR using relative gene expression (ΔΔC_T_ Method), the statistical significance were assessed based on ANOVA using ΔCt (*55*). Briefly, total RNA was extracted from PDO using the Norgen single cell RNA purification kit (Norgen, 51800). The TagMan primers and probes were obtained from Thermo Fisher, and ACTB was leveraged as the endogenous control (KLF14: Hs00370951_s1; EGR2: Hs00370951_s1; EPHA4: Hs00953178_m1; ACTB: HS01060665-g1). Organoid growth images were taken by the Incucyte® system.

#### Compound Sensitivity Testing

PDO with EPHA4 shRNAs or scramble control were seeded into 96 well plates at 3000 viable cells per cell with 10 µL Matrigel. Then 100 µL CRC culture medium containing RealTime-Glo™ MT Cell Viability Assay (Promega G9711) and compounds of 10-point dose curve along with DMSO control was added. The cell viability at 0 hours for each well was measured by RealTime-Glo™ MT Cell Viability Assay for growth rate correction. The organoids were treated (drug or DMSO) for 5 days prior to assessing for viabilities. The PDO viabilities were determined by CellTiter-Glo® 3D Cell Viability Assay (Promega, G9683). Briefly, 100 µL CellTiter-Glo® 3D was added to each well, then the plate was shaken for 5 min followed by a 20 min incubation at room temperature. The luminescence was measured by the plate reader (Varioskan™ LUX). Each dose point was normalized to the DMSO control for relative viability. The growth-rate dose response curve fit a 3-parameter sigmoidal curve using GraphPad Prism 9. The compounds used for the testing were 5-Fluorouracil (Millipore Sigma, F6627-5G), Mirdametinib (PD0325901) (Millipore Sigma, PZ0162-25MG), Pemigatinib (Selleckchem, Catalog No.S0088), and Erdafitinib (Selleckchem, Catalog No.S8401). The high throughput drug screen was performed by Duke Functional Genomics Core. Briefly, the NIH approved oncology drug set IX (plate 4891 and 4892, including 147 compounds) was used for the screen. The EPHA4 overexpression (OE) PDO were transduced with EPHA4 ORF lentivirus (EPHA4_OHu24718C_pGenlenti, GenScript) (Figure S12D). The EPHA4 shRNA1 lentivirus were used for EPHA4 knocking down PDO. The PDO were exposed to the compounds at 1 µM (N=3 for each compound & each PDO condition) for three days. Then the viabilities were determined by CellTiter-Glo® 3D Cell Viability Assay. The luminescence for each drug was normalized to the DMSO treated PDO. The heatmap was generated by the R package ComplexHeatmap with the K-nearest-neighbor clustering km=4.

#### In Vivo PDOX Growth Measurements

PDO with distinct types of shRNAs were harvested, counted, and resuspended in 50% Matrigel PBS solution at a concentration of 10^7^ cells/mL. PDO solution (100 µL, 1 million cells) was injected s.c. into the flanks of an NSG mouse using a 23-G needle. The tumor length and width were measured with a 0.01-mm vernier caliper every three days. The tumor volume (V) was determined as *V* = *width*^2^ × *length* ÷ 2. At the end of the study, the tumors were harvested and weighed. Five mice for each type of PDO were used, same as the scramble control group.

#### ATAC-seq and mRNA-seq Library Preparation and Sequencing

We removed the mouse cell components from the xenograft samples and dead cells from all DMSO preserved samples before preparing the sequencing libraries. Briefly, cells were recovered from the DMSO cryopreservation, then were washed with PBS and spun down. The supernatant was removed and resuspended in 50 µL mouse depletion cocktail (for the xenografts samples, Miltenyi Biotech, #130-104-694) + 100 µL dead cell removal microbeads (Miltenyi Biotech, #130-090-101) + 60 µL binding buffer and incubated 15 min at room temperature. The mixture was passed through an LS column (Miltenyi Biotech, #130-042-401), and the LS column was washed three times with 2 mL binding buffer. The live human cells were in the flow-through and ready for the next step. The Omni-ATAC protocol (*23*) was utilized for ATAC library preparations. The nuclei were extracted from samples (PT, PDO, PDX, and PDOX). For each sample type in one PT-PDO-PDX/PDOX set (e.g., PDOX19-106), three replicates were included (Xenografts were from three mice). Intact nuclei (5 × 10^4^) were used for each replicate. Transposase, digitonin, and Tween-20 were added, and the mixture was incubated at 37 °C for 30 minutes. The transposed fragments were amplified. The cycle numbers of amplifications were determined by qPCR for each sample. The libraries that passed QC were sequenced by Illumina HiSeq 4000 PE150 bp. The RNA was extracted using the Norgen single cell RNA purification kit (Norgen, 51800). The mRNA-seq libraries were made by Novogene and sequenced by Illumina HiSeq 4000 PE150 bp. Chromium Next GEM single cell Multiome kit (10X Genomics) was utilized for single cell multiome ATAC + gene expression library preparation. Briefly, the 19-187 PT-PDMC set with patient, PDO, PDOX, and PDX samples was used for 10X single cell sequencing. 8000 nuclei from each sample were extracted and transposed. Then, the GEMs were generated using the Chromium Next GEM Chip J and barcodes were added. After cleanup and pre-amplification, the single cell ATAC libraries and the single cell gene expression libraries were constructed following the 10X protocol (CG000338 Rev B). The libraries were sequenced using NovaSeq 6000 S1 PE100 bp.

#### ATAC-seq and mRNA-seq Data Processing

ATAC-seq data was processed using the pipeline developed for ENCODE (https://github.com/ENCODE-DCC/atac-seq-pipeline) to perform quality control, filtering of low-quality reads and PCR duplicates, analysis of reproducibility, reference genome alignment, peaks calling, and fold-enrichment or p-value signal tracks generation. Reads were aligned to reference genome hg19 using Bowtie2 (*56*). Duplicate and mitochondrial reads were removed, and peaks were called by MACS2 (*57*).

RNA-seq raw sequencing reads were quality checked using Fastqc and summarized with MultiQC (*58*). Transcripts were aligned to reference genome hg19 using Hisat2 and quantified by HTSeq (*59, 60*). The mutation information was obtained from the mRNA-seq data. Briefly, The SNPs and Indels were called from raw RNA-seq reads following the GATK Best Practices RNA-seq workflow (*61*). The final annotation step was performed by ANNOVAR (*62*). The oncoplot was generated by the R package maftools (*63*).

#### Differential Peak Calling and Annotation

Differential peaks were identified using DiffBind with a minimum of three biological replicates (*24*). The fold-changes of peaks were evaluated by the DESEQ2 method in the DiffBind package. Peaks with p-value <= 0.05 and |log2FC| >= 1 were considered differentially accessible. Peaks annotation was performed using the findMotifsGenome.pl command in the HOMER2 software based on the nearest TSS (*64*). The R package ChIPseeker was also used to annotate peaks and visualization (*65*). Gene Ontology was performed by the R package clusterProfiler (*66*). For mRNA-seq, Deseq2 was used for differential expression analyses. Genes differentially expressed were classified by p-value <= 0.05 and log2FC >= 1 (*67*).

### ATAC Track Visualization and Correlation

Bam files from each ATAC library were used as input for deeptool2 bamCoverage command to generate the bigwig files (*68*). The bigwig files were visualized by the R package Gviz. The correlations between different bam files were performed using the deeptool2 multiBamsummary command and the resulting matrix was plotted by the plotCorrelation --whatToPlot scatterplot command.

### Sequencing Data Visualization

Circos plots were generated based on the differential or consensus peaks generated by Diffbind using the R package circlize (*69*). Briefly, circos.genomicDensity function was used to visualize the genomic density of the genome.

Heatmaps, PCA plots, and MA plots were generated using the built-in Diffbind methods. The heatmap for clustering was generated based on the 1000 peaks with the highest variance. The PCA plot was generated based on the DBA_SCORE_RPKM of each replicate. The heatmap for chromatin accessibility around a TSS was generated using deepTools (*68*).

### ATAC Differential Occupancy and Pathway Analysis

The consensus peak set for the PDMC models and patient ATAC-seq results were generated by the R package Diffbind. In short, a count matrix was generated for the combined consensus peak set, and the counts were normalized with DBA_NORM_NATIVE and DBA_LIBSIZE_PEAKREADS options. Three contrasts: 1) PDO vs. Patient, 2) PDOX vs. Patient and 3) PDX vs. Patient were generated and used as input for dba.analyze with method=DBA_DESEQ2. The resulting DE peaks from all three contrasts were summarized into a data frame, and K-means clustering with centers=4 was performed to cluster the peaks into 4 different clusters. The clustered peaks were then visualized by R package ComplexHeatmap (*70*). The shared gain and loss of peaks from different comparisons were the subjected to GREAT for pathway analysis (*71*).

### 10X Single-cell Analysis

The Illumina sequenced 10x single-cell multiome libraries were subjected to the 10x Cell-Ranger-Arc pipeline with default steps. Briefly, sequence data was demultiplexed into single-cell ATAC results and RNA results by running the cellranger-arc mkfastq command. Single cell feature counts for both ATAC and RNA results were then generated by the cellranger-arc count command, with GRCh38 as a reference genome. Samples sequenced from different sequencing lanes were aggregated by the cellranger-arc aggr command. The resulting count matrices for each PDMC and patient model were then treated as inputs for downstream analysis. The expression matrices were also subjected to SoupX to remove any predicted ambient RNA contamination with default parameters(*72*). Seurat objects were then created from the resulting Gene expression matrix for further aggregation and population identification(*73, 74*). The filtering scheme are nCount_ATAC < 100000 & nCount_ATAC > 1000 (200 for PT); nCount_RNA < 100000 (30000 for PT) & nCount_RNA > 1000 (500 for PDX, 200 for PT); nFeature_RNA > 1000 (500 for PDX, 100 for PT) & nFeature_RNA<10000; nucleosome_signal < 2; TSS.enrichment > 1; percent.mt < 30 (25 for PDX, 10 for PT).

### Transcription Factor Analysis

BaGFoot Analysis was performed following the previously described methods (*29*). The hg19 genome was used as the reference. Aligned and filtered ATAC-seq peaks from each biological replicate were pooled to create a consensus file for each condition and used for pairwise BaGFoot analysis. The outer polygon fence was created by inflating the bag geometrically by a factor of 1.5.

Footprint analysis was performed using the TOBIAS BINDetect tool (*33*). The meme file was downloaded from the HOCOMOCO database. Correction of Tn5 insertion bias, calculation of footprint scores within regulatory regions, estimation of bound/unbound transcription factor binding sites, and visualization of footprints within and across different conditions were performed using default parameters.

### Identification of Downstream Genes for KLF14 and EGR2

In the ATAC-seq and RNA-seq, DE analysis for PDOX vs. PDO, the differentially enriched peaks (ATAC-seq) are annotated to the nearest transcription start site and denoted as ATAC-seq DE genes. The PDOX ATAC-seq and RNA-seq DE genes are thereby inner joined to find the PDOX ATAC+RNA-seq enriched gene set. The downstream genes regulated by KLF14 and EGR2 were identified by using the R package tftargets. Supplemental Table 5 was generated by taking the intersect of the KLF14 and EGR2 downstream genes from the Marbach 2016 dataset (*75*) with the PDOX ATAC+RNA-seq enriched gene set.

### Statistical Analysis

DiffBind and DESeq2 were used for the differential analyses of ATAC-seq data, and DESeq2 was used for mRNA-seq. Animal data were expressed as mean ± standard error of the mean (SEM) of at least three biological replicates. The number of biological replicates was denoted in the figure legends. Paired Student’s t-test or ANOVA were used for comparisons, with p-value <= 0.05 considered to be significant. Mice were randomly allocated to experimental groups.

## Supplemental Materials

**Figure S1 Histological staining of patient and PDMC**.

**Figure S2 Pairwise ATAC-seq correlation between different replicates of patient samples and models**.

**Figure S3 Circos plot based on ATAC-seq data**.

**Figure S4 Supplemental principal component analysis and correlation analysis**.

**Figure S5 Supplemental MA plots based on ATAC-seq data**.

**Figure S6 The pathway analysis**.

**Figure S7 Supplemental single cell multiome analysis**.

**Figure S8 Pathway Enrichment Analysis of PDOX vs. PDO**.

**Figure S9 Supplemental footprint analysis**.

**Figure S10 BaGFoot analysis for individual patient sets**.

**Figure S11 Supplemental KLF14 and EGR2 *in vivo* validations. Figure S12 EPHA4 related drug sensitivity tests**.

**Supplemental Table 1 Patient Information**

**Supplemental Table 2 Sequencing Summary**

**Supplemental Table 3 Differential Peak Number**

**Supplemental Table 4 Transcription Factors Analysis**

**Supplemental Table 5 Joined KLF14 & EGR2 PDOX Downstream Genes**

## Notes

### Competing Interest Statement

The authors have declared no competing interest.

