## Supplemental Figures S1_S12 for "Chromatin Remodeling in Patient-Derived Colorectal Cancer Models"

Figure S1

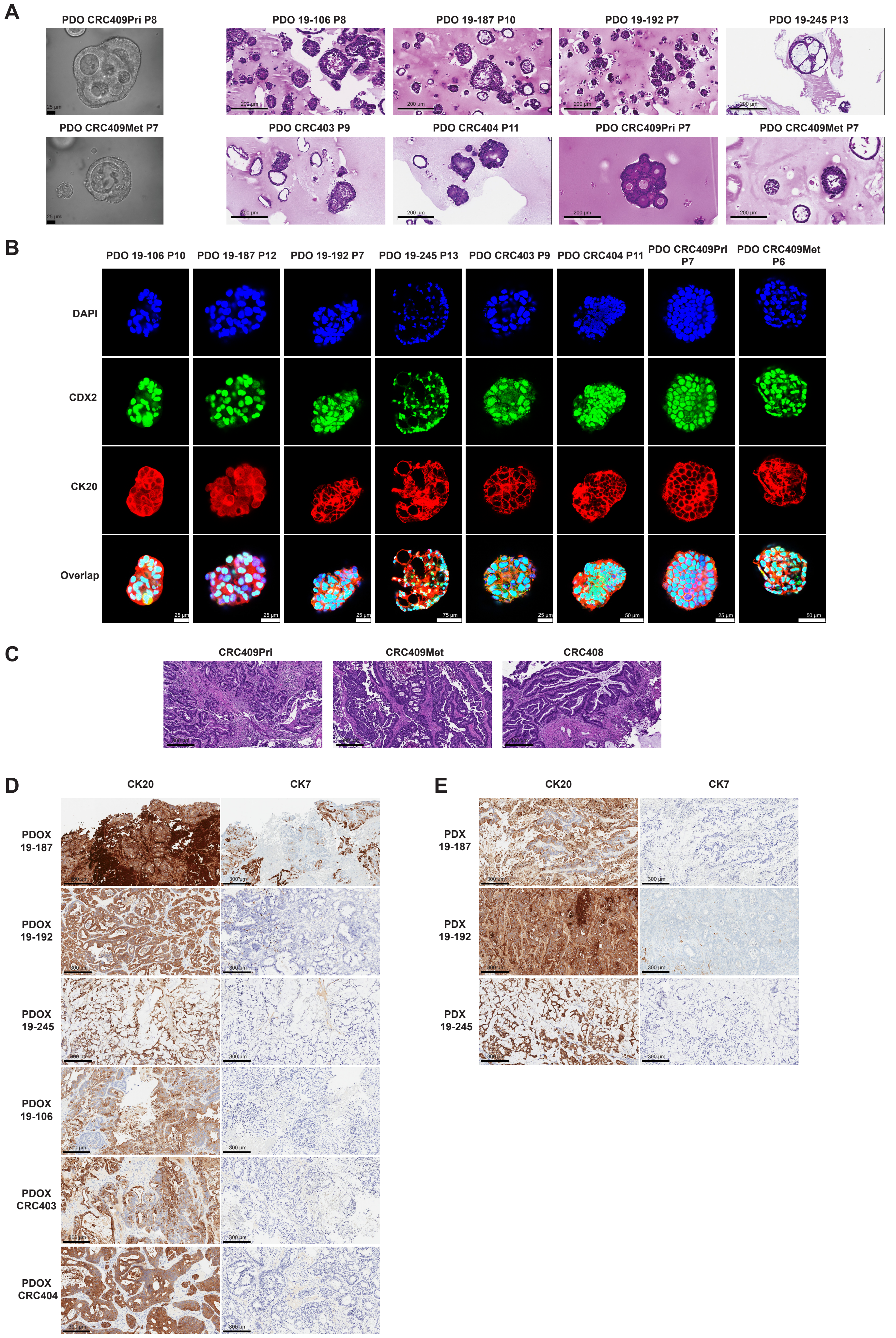
A
PDO CRC409Pri P8
PDO 19-106 P8
PDO 19-187 P10
PDO 19-192 P7
PDO 19-245 P13
PDO CRC409Met P7
PDO CRC403 P9
PDO CRC404 P11
PDO CRC409Pri P7
PDO CRC409Met P7
B
PDO 19-106 P10
PDO 19-187 P12
PDO 19-192 P7
PDO 19-245 P13
PDO CRC403 P9
PDO CRC404 P11
PDO CRC409Pri P7
PDO CRC409Met P6
DAPI
CDX2
CK20
Overlap
C
CRC409Pri
CRC409Met
CRC408
D
CK20
CK7
PDOX 19-187
PDOX 19-192
PDOX 19-245
PDOX 19-106
PDOX CRC403
PDOX CRC404
E
CK20
CK7
PDX 19-187
PDX 19-192
PDX 19-245

**Figure S1 Histological staining of patient and PDMC.**

**(A)** Left panel: PDO bright-field images (40X objective). Scale bars, 25 µm. Right panel: H&E-staining of paraffin-embedded PDO. Scale bars, 200 µm.

**(B)** High resolution whole-mount confocal image of PDO immunolabeled for CDX2 (green) and CK20 (red). DAPI is labeled in blue. The overlap images are also shown in Figure 1C. Scale bars, 75 µm for PDO 19-245, 50 µm for PDO CRC404, and 25 µm for the rest.

**(C)** H&E-staining of patient samples. Scale bars, 300 µm.

**(D)** IHC staining for CK20 (left) and CK7 (right) of PDOX. Scale bars, 300 µm.

**(E)** IHC staining for CK20 (left) and CK7 (right) of PDX. Scale bars, 300 µm.

Figure S2

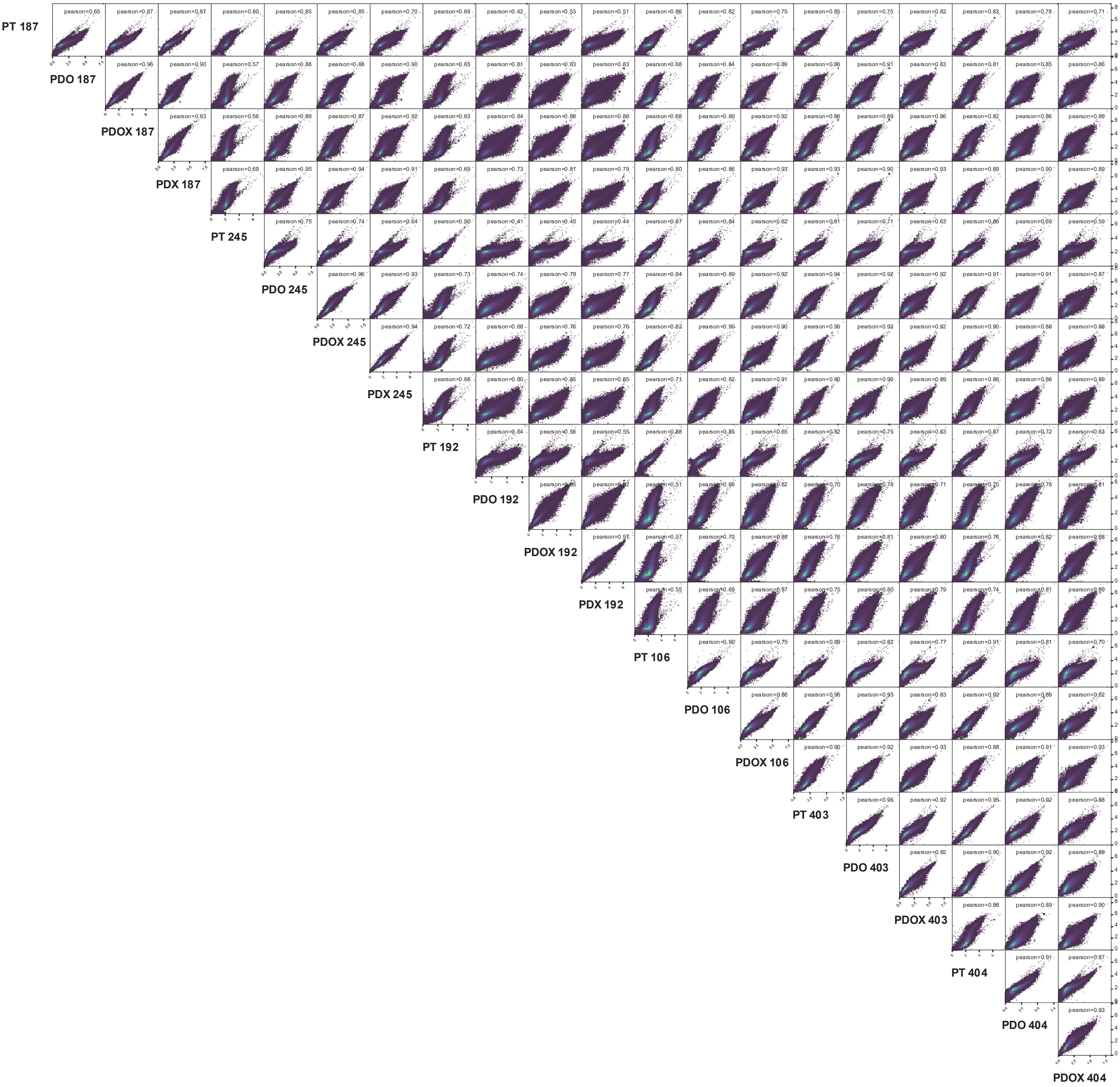
PT 187
PDO 187
PDOX 187
PDX 187
PT 245
PDO 245
PDOX 245
PDX 245
PT 192
PDO 192
PDOX 192
PDX 192
PT 106
PDO 106
PDOX 106
PT 403
PDO 403
PDOX 403
PT 404
PDO 404
PDOX 404

**Figure S2** Pairwise ATAC-seq correlation between different replicates of patient samples and models based on normalized read counts of detected peaks. Pearson correlation coefficient is presented for each comparison.

Figure S3

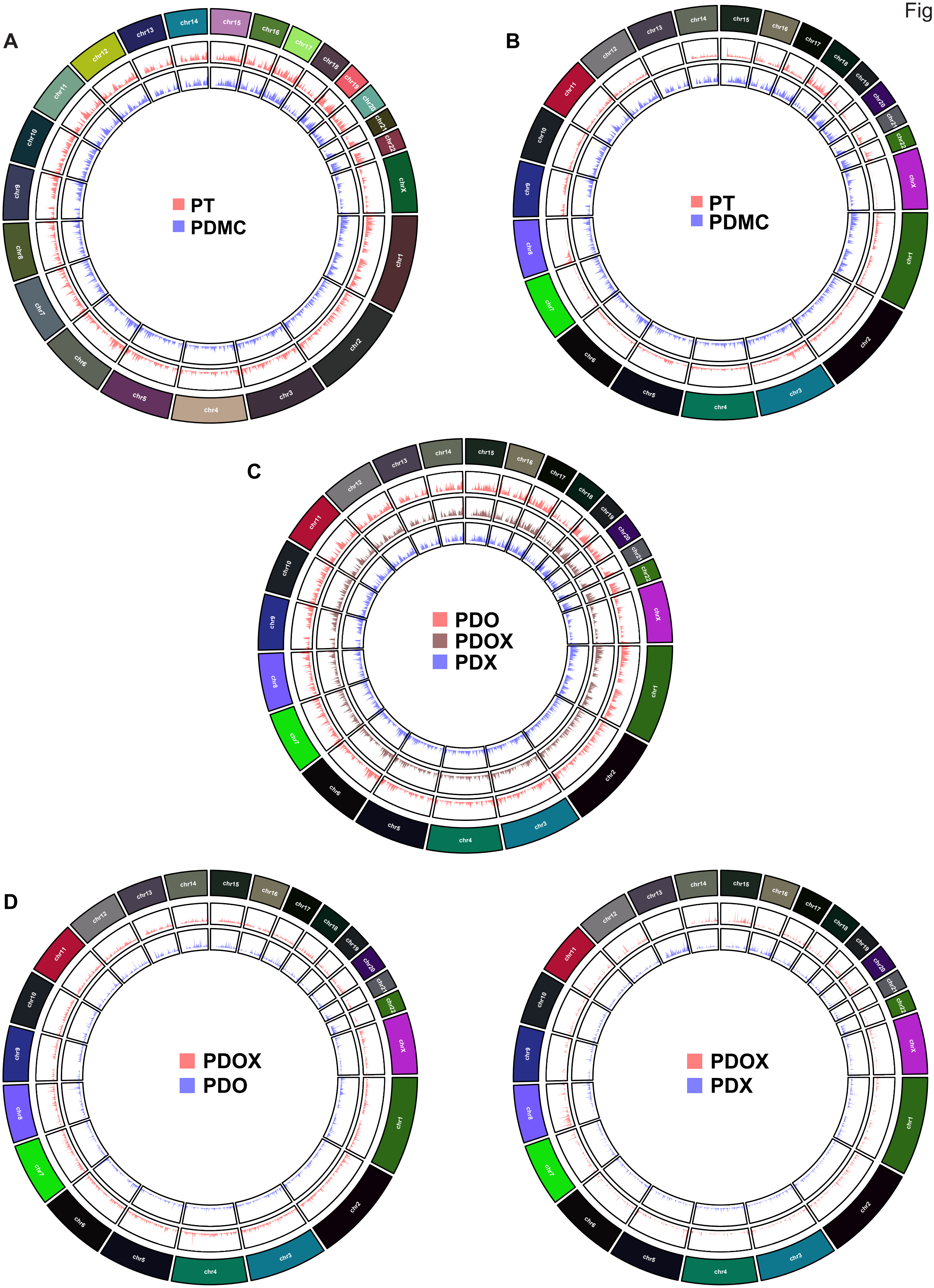
A
PT
PDMC
B
PT
PDMC
C
PDO
PDOX
PDX
D
PDOX
PDO
PDOX
PDX
chr1
chr2
chr3
chr4
chr5
chr6
chr7
chr8
chr9
chr10
chr11
chr12
chr13
chr14
chr15
chr16
chr17
chr18
chr19
chr20
chr21
chr22
chrX

**Figure S3 Circos plot based on ATAC-seq data.**

**(A)** Circos plot showing the global chromatin accessibility based on ATAC-seq. The accumulated ATAC-seq peak density was presented in peak positions within the chromosomes. The peaks of PT are shown in red, and the peaks of PDMC are blue.

**(B)** Circos plot showing the DE peaks density of the comparison of PT vs. PDMC ($|log2(FC)|>1$, $p<0.05$).

**(C)** Circos plot showing the global chromatin accessibilities of PDO, PDOX, and PDX.

**(D)** Circos plot showing the DE peaks density of the comparison of PDOX vs. PDO and PDOX vs. PDX ($|log2(FC)|>1$, $p<0.05$).

Figure S4

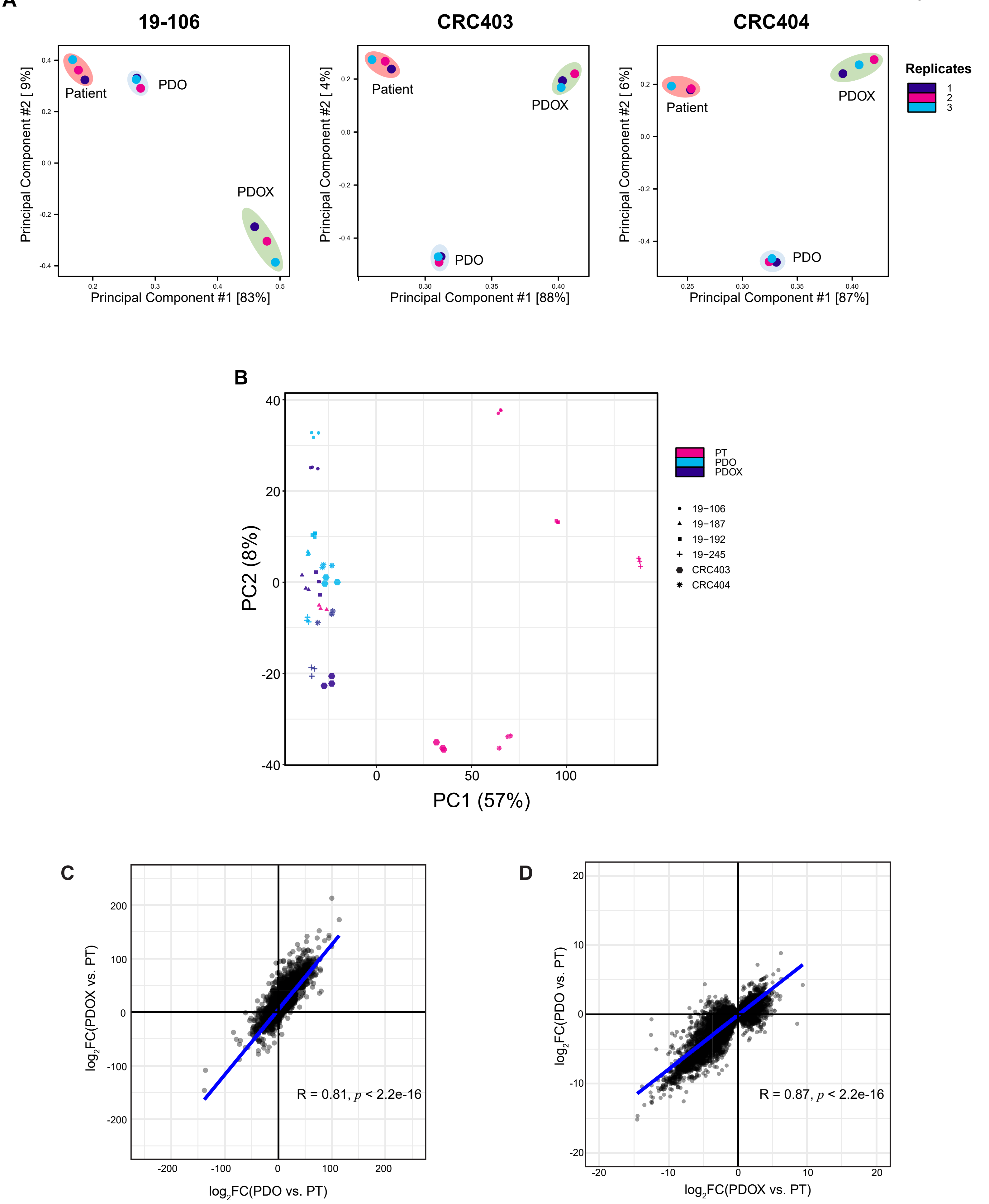

A
19-106
CRC403
CRC404
Patient
PDO
PDOX
Principal Component #2 [ 9%]
Principal Component #1 [83%]
Principal Component #2 [ 4%]
Principal Component #1 [88%]
Principal Component #2 [ 6%]
Principal Component #1 [87%]
Replicates
1
2
3
B
PC2 (8%)
PC1 (57%)
PT
PDO
PDOX
19−106
19−187
19−192
19−245
CRC403
CRC404
C
log2FC(PDOX vs. PT)
log2FC(PDO vs. PT)
R = 0.81, p < 2.2e-16
D
log2FC(PDO vs. PT)
log2FC(PDOX vs. PT)
R = 0.87, p < 2.2e-16

**Figure S4 Supplemental principal component analysis and correlation analysis.**

**(A)** Principal component analysis of individual PT-PDMC sets (19-106, CRC403, and CRC404) from ATAC-seq data based on the DiffBind score DBA_SCORE_RPKM. Patient samples are circled in red, PDO in blue, and PDOX in light green. Each type of model has three replicates.

**(B)** Principal component analysis of the pooled PT-PDO-PDOX sets based on mRNA-seq data. Magenta indicates patient samples, cyan indicates PDO, and dark violet indicates PDOX. Different shapes of dots denote patient IDs.

**(C)** Correlation analysis of PDOX vs. PT and PDO vs. PT based on the differential enrichment (log2FC) ATAC-seq. Pearson correlation R value is 0.81.

**(D)** Correlation analysis of PDOX vs. PT and PDO vs. PT based on the differential expression (log2FC) in mRNA-seq. Pearson correlation R value is 0.87.

Figure S5

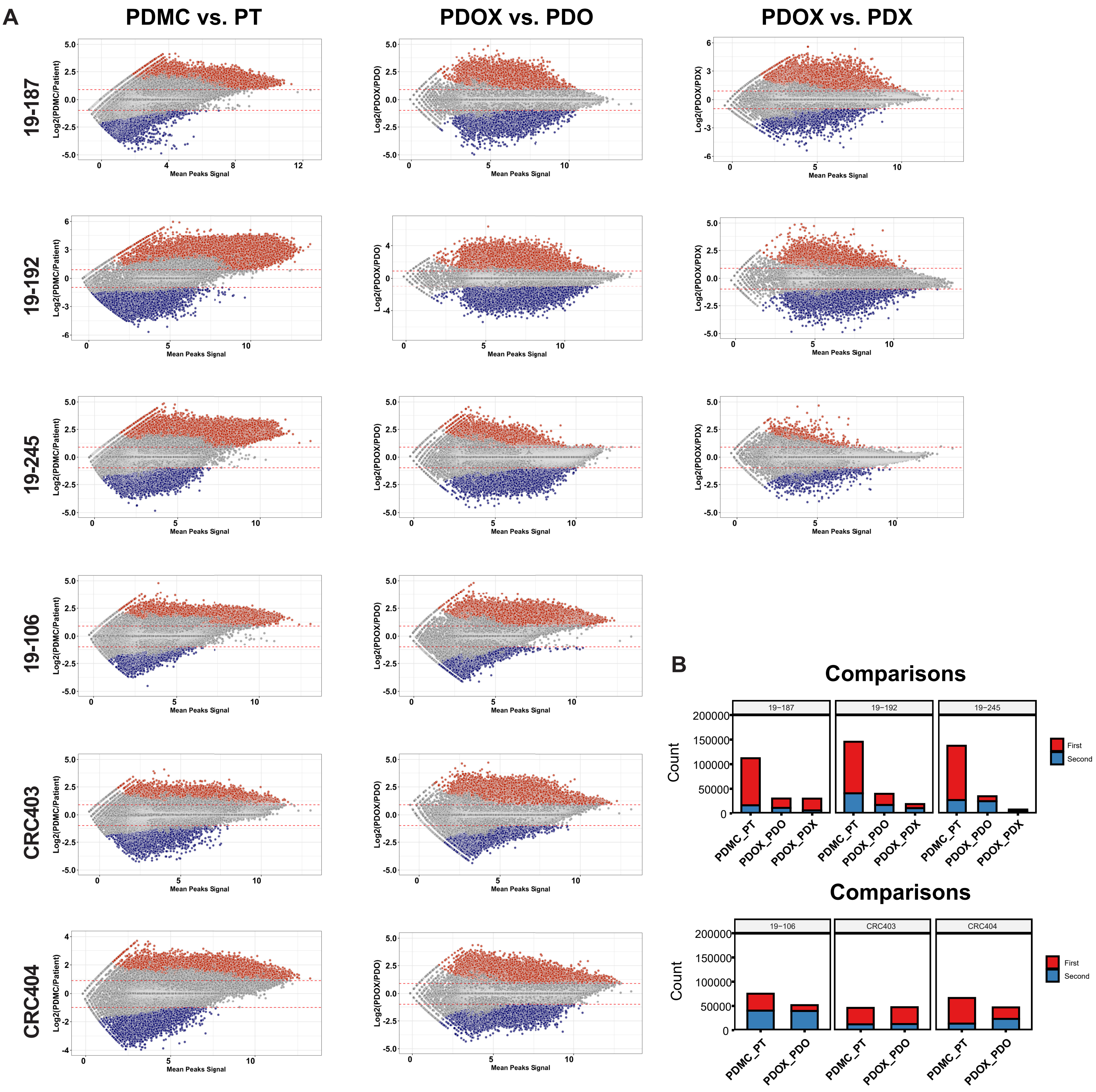
A
PDMC vs. PT
PDOX vs. PDO
PDOX vs. PDX
19-187
19-192
19-245
19-106
CRC403
CRC404
Log2(PDMC/Patient)
Log2(PDOX/PDO)
Log2(PDOX/PDX)
Mean Peaks Signal
B
Comparisons
Count
First
Second
19−187
19−192
19−245
19−106
CRC403
CRC404
PDMC_PT
PDOX_PDO
PDOX_PDX

**Figure S5 Supplemental MA plots based on ATAC-seq data.**

**(A)** ATAC-seq differential analysis of individual PT-PDMC sets. MA plots demonstrating the differential enriched peaks in each comparison (|log2(FC)|>1, $p<0.05$). **(B)** The bar plots report the number of counts of the differentially enriched peaks.

Figure S6

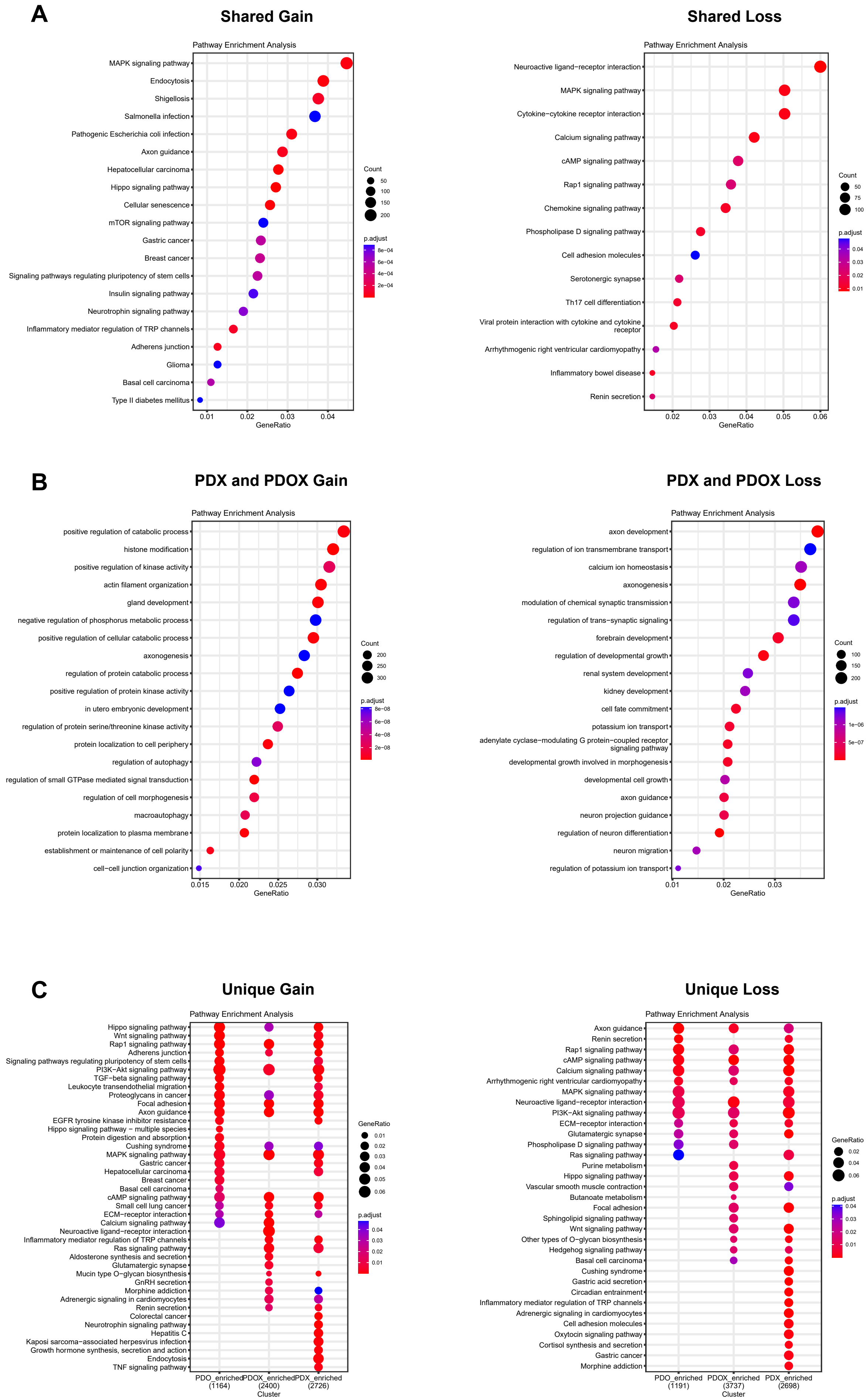

A
Shared Gain
Pathway Enrichment Analysis
MAPK signaling pathway
Endocytosis
Shigellosis
Salmonella infection
Pathogenic Escherichia coli infection
Axon guidance
Hepatocellular carcinoma
Hippo signaling pathway
Cellular senescence
mTOR signaling pathway
Gastric cancer
Breast cancer
Signaling pathways regulating pluripotency of stem cells
Insulin signaling pathway
Neurotrophin signaling pathway
Inflammatory mediator regulation of TRP channels
Adherens junction
Glioma
Basal cell carcinoma
Type II diabetes mellitus
0.01 0.02 0.03 0.04
GeneRatio
Count
50
100
150
200
p.adjust
8e−04
6e−04
4e−04
2e−04
Shared Loss
Pathway Enrichment Analysis
Neuroactive ligand−receptor interaction
MAPK signaling pathway
Cytokine−cytokine receptor interaction
Calcium signaling pathway
cAMP signaling pathway
Rap1 signaling pathway
Chemokine signaling pathway
Phospholipase D signaling pathway
Cell adhesion molecules
Serotonergic synapse
Th17 cell differentiation
Viral protein interaction with cytokine and cytokine receptor
Arrhythmogenic right ventricular cardiomyopathy
Inflammatory bowel disease
Renin secretion
0.02 0.03 0.04 0.05 0.06
GeneRatio
Count
50
75
100
p.adjust
0.04
0.03
0.02
0.01
B
PDX and PDOX Gain
Pathway Enrichment Analysis
positive regulation of catabolic process
histone modification
positive regulation of kinase activity
actin filament organization
gland development
negative regulation of phosphorus metabolic process
positive regulation of cellular catabolic process
axonogenesis
regulation of protein catabolic process
positive regulation of protein kinase activity
in utero embryonic development
regulation of protein serine/threonine kinase activity
protein localization to cell periphery
regulation of autophagy
regulation of small GTPase mediated signal transduction
regulation of cell morphogenesis
macroautophagy
protein localization to plasma membrane
establishment or maintenance of cell polarity
cell−cell junction organization
0.015 0.020 0.025 0.030
GeneRatio
Count
200
250
300
p.adjust
8e−08
6e−08
4e−08
2e−08
PDX and PDOX Loss
Pathway Enrichment Analysis
axon development
regulation of ion transmembrane transport
calcium ion homeostasis
axonogenesis
modulation of chemical synaptic transmission
regulation of trans−synaptic signaling
forebrain development
regulation of developmental growth
renal system development
kidney development
cell fate commitment
potassium ion transport
adenylate cyclase−modulating G protein−coupled receptor signaling pathway
developmental growth involved in morphogenesis
developmental cell growth
axon guidance
neuron projection guidance
regulation of neuron differentiation
neuron migration
regulation of potassium ion transport
0.01 0.02 0.03
GeneRatio
Count
100
150
200
p.adjust
1e−06
5e−07
C
Unique Gain
Pathway Enrichment Analysis
Hippo signaling pathway
Wnt signaling pathway
Rap1 signaling pathway
Adherens junction
Signaling pathways regulating pluripotency of stem cells
PI3K−Akt signaling pathway
TGF−beta signaling pathway
Leukocyte transendothelial migration
Proteoglycans in cancer
Focal adhesion
Axon guidance
EGFR tyrosine kinase inhibitor resistance
Hippo signaling pathway − multiple species
Protein digestion and absorption
Cushing syndrome
MAPK signaling pathway
Gastric cancer
Hepatocellular carcinoma
Breast cancer
Basal cell carcinoma
cAMP signaling pathway
Small cell lung cancer
ECM−receptor interaction
Calcium signaling pathway
Neuroactive ligand−receptor interaction
Inflammatory mediator regulation of TRP channels
Ras signaling pathway
Aldosterone synthesis and secretion
Glutamatergic synapse
Mucin type O−glycan biosynthesis
GnRH secretion
Morphine addiction
Adrenergic signaling in cardiomyocytes
Renin secretion
Colorectal cancer
Neurotrophin signaling pathway
Hepatitis C
Kaposi sarcoma−associated herpesvirus infection
Growth hormone synthesis, secretion and action
Endocytosis
TNF signaling pathway
PDO_enriched (1164)
PDOX_enriched (2400)
PDX_enriched (2726)
Cluster
GeneRatio
0.01
0.02
0.03
0.04
0.05
0.06
p.adjust
0.04
0.03
0.02
0.01
Unique Loss
Pathway Enrichment Analysis
Axon guidance
Renin secretion
Rap1 signaling pathway
cAMP signaling pathway
Calcium signaling pathway
Arrhythmogenic right ventricular cardiomyopathy
MAPK signaling pathway
Neuroactive ligand−receptor interaction
PI3K−Akt signaling pathway
ECM−receptor interaction
Glutamatergic synapse
Phospholipase D signaling pathway
Ras signaling pathway
Purine metabolism
Hippo signaling pathway
Vascular smooth muscle contraction
Butanoate metabolism
Focal adhesion
Sphingolipid signaling pathway
Wnt signaling pathway
Other types of O−glycan biosynthesis
Hedgehog signaling pathway
Basal cell carcinoma
Cushing syndrome
Gastric acid secretion
Circadian entrainment
Inflammatory mediator regulation of TRP channels
Adrenergic signaling in cardiomyocytes
Cell adhesion molecules
Oxytocin signaling pathway
Cortisol synthesis and secretion
Gastric cancer
Morphine addiction
PDO_enriched (1191)
PDOX_enriched (3737)
PDX_enriched (2698)
Cluster
GeneRatio
0.02
0.04
0.06
p.adjust
0.04
0.03
0.02
0.01

**Figure S6 The pathway analysis.**

**(A)** KEGG pathways enriched in the shared gain or loss peaks defined in Figure 2I. **(B)** GeneOntology biological processes enriched in the shared gain/loss peaks in PDX and PDOX as defined in Figure 2I. **(C)** KEGG pathways enriched in the unique gain/loss peaks for PDO, PDOX and PDX as defined in Figure 2I.

Figure S7

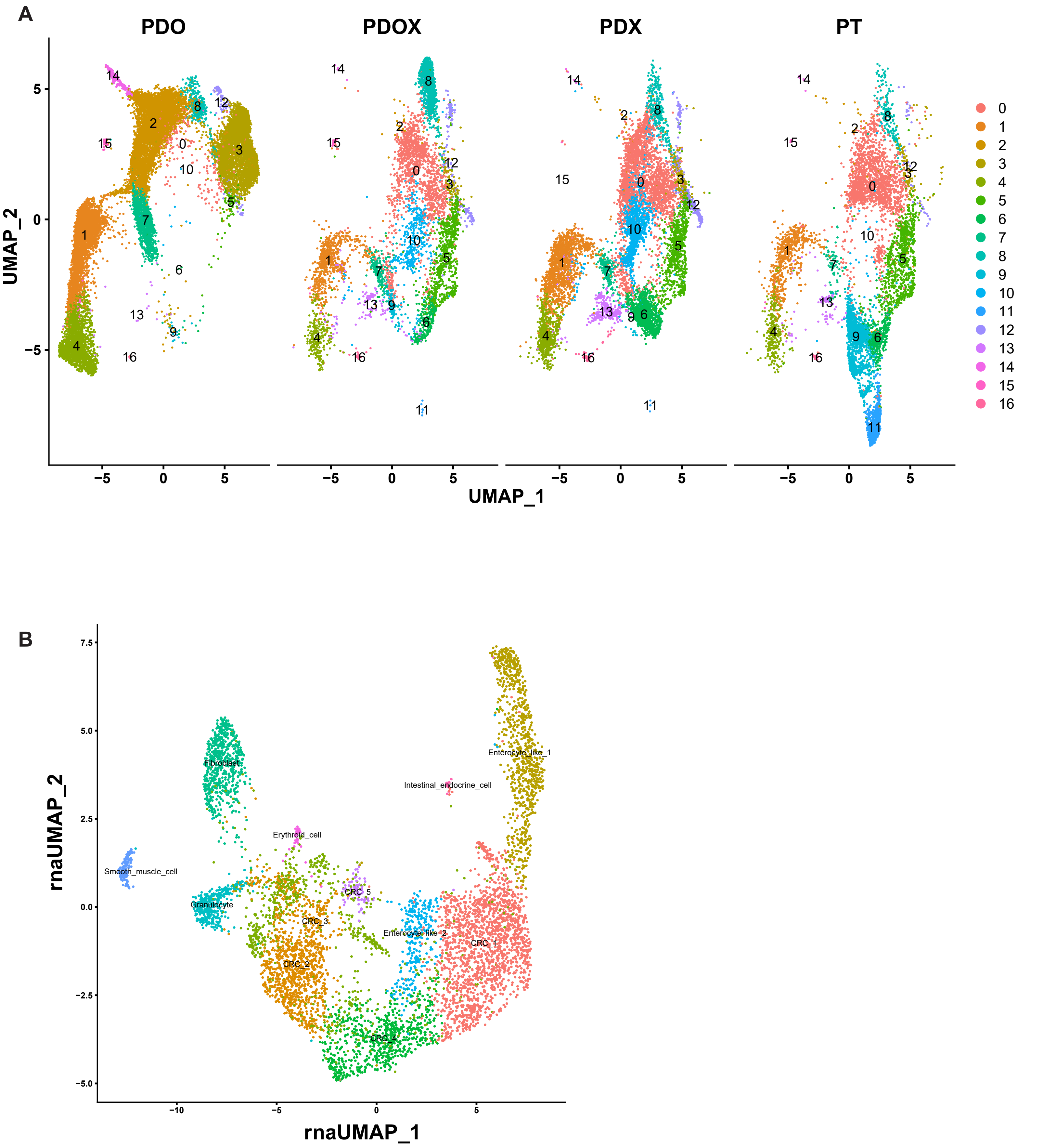
A
PDO
PDOX
PDX
PT
UMAP_2
UMAP_1
B
rnaUMAP_2
rnaUMAP_1
Fibroblast
Smooth_muscle_cell
Granulocyte
Erythroid_cell
Intestinal_endocrine_cell
Enterocyte_like_1
Enterocyte_like_2
CRC_1
CRC_2
CRC_3
CRC_4
CRC_5

**Figure S7 Supplemental single cell multiome analysis.**

**(A)** Integrated UMAP of PDO, PDOX, PDX, and PT samples from 19-187 set, clustering based on gene expressions.

**(B)** Visualizations of the cell clusters based on the gene expression in the PT sample only, showing the cell types of non-malignant /CRC components.

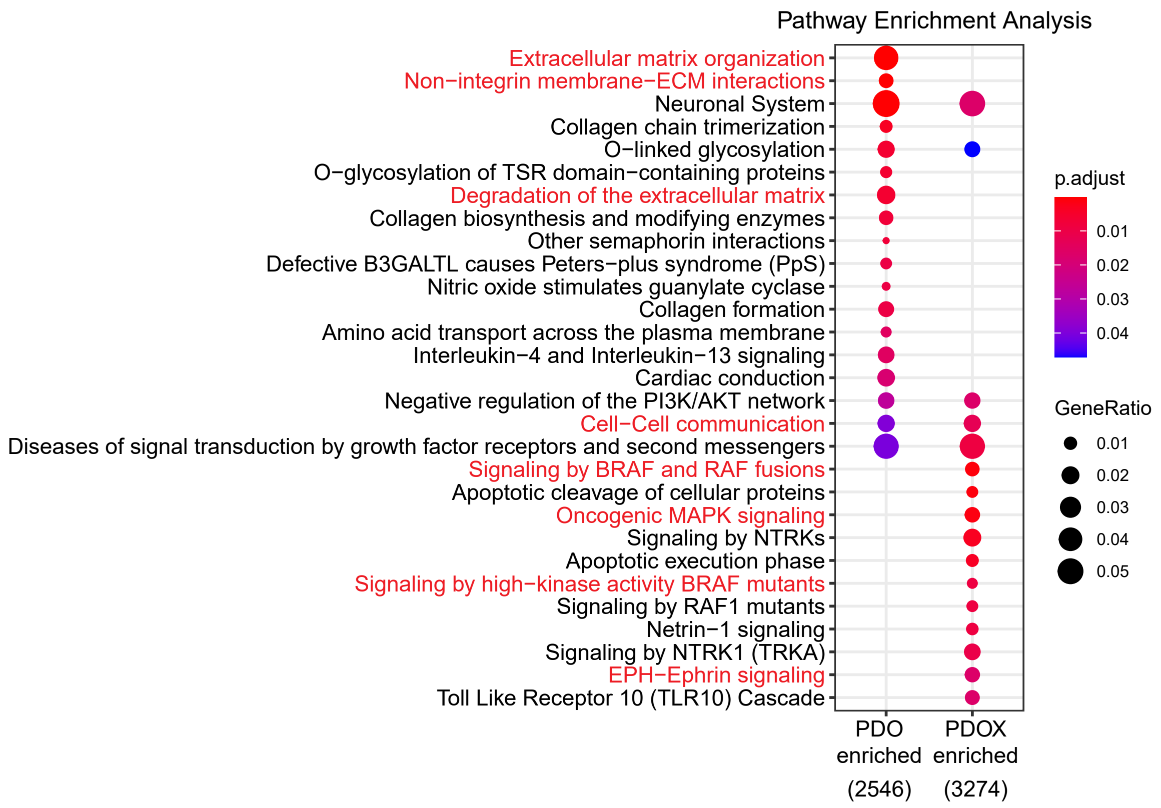
Pathway Enrichment Analysis
Extracellular matrix organization
Non−integrin membrane−ECM interactions
Neuronal System
Collagen chain trimerization
O−linked glycosylation
O−glycosylation of TSR domain−containing proteins
Degradation of the extracellular matrix
Collagen biosynthesis and modifying enzymes
Other semaphorin interactions
Defective B3GALTL causes Peters−plus syndrome (PpS)
Nitric oxide stimulates guanylate cyclase
Collagen formation
Amino acid transport across the plasma membrane
Interleukin−4 and Interleukin−13 signaling
Cardiac conduction
Negative regulation of the PI3K/AKT network
Cell−Cell communication
Diseases of signal transduction by growth factor receptors and second messengers
Signaling by BRAF and RAF fusions
Apoptotic cleavage of cellular proteins
Oncogenic MAPK signaling
Signaling by NTRKs
Apoptotic execution phase
Signaling by high−kinase activity BRAF mutants
Signaling by RAF1 mutants
Netrin−1 signaling
Signaling by NTRK1 (TRKA)
EPH−Ephrin signaling
Toll Like Receptor 10 (TLR10) Cascade
PDO enriched (2546)
PDOX enriched (3274)
p.adjust
0.01
0.02
0.03
0.04
GeneRatio
0.01
0.02
0.03
0.04
0.05

**Figure S8** Pathway Enrichment Analysis of PDOX vs. PDO.

KEGG pathways enrichment analysis based on the differentially enriched peaks in the comparison of PDOX vs. PDO.

Figure S9

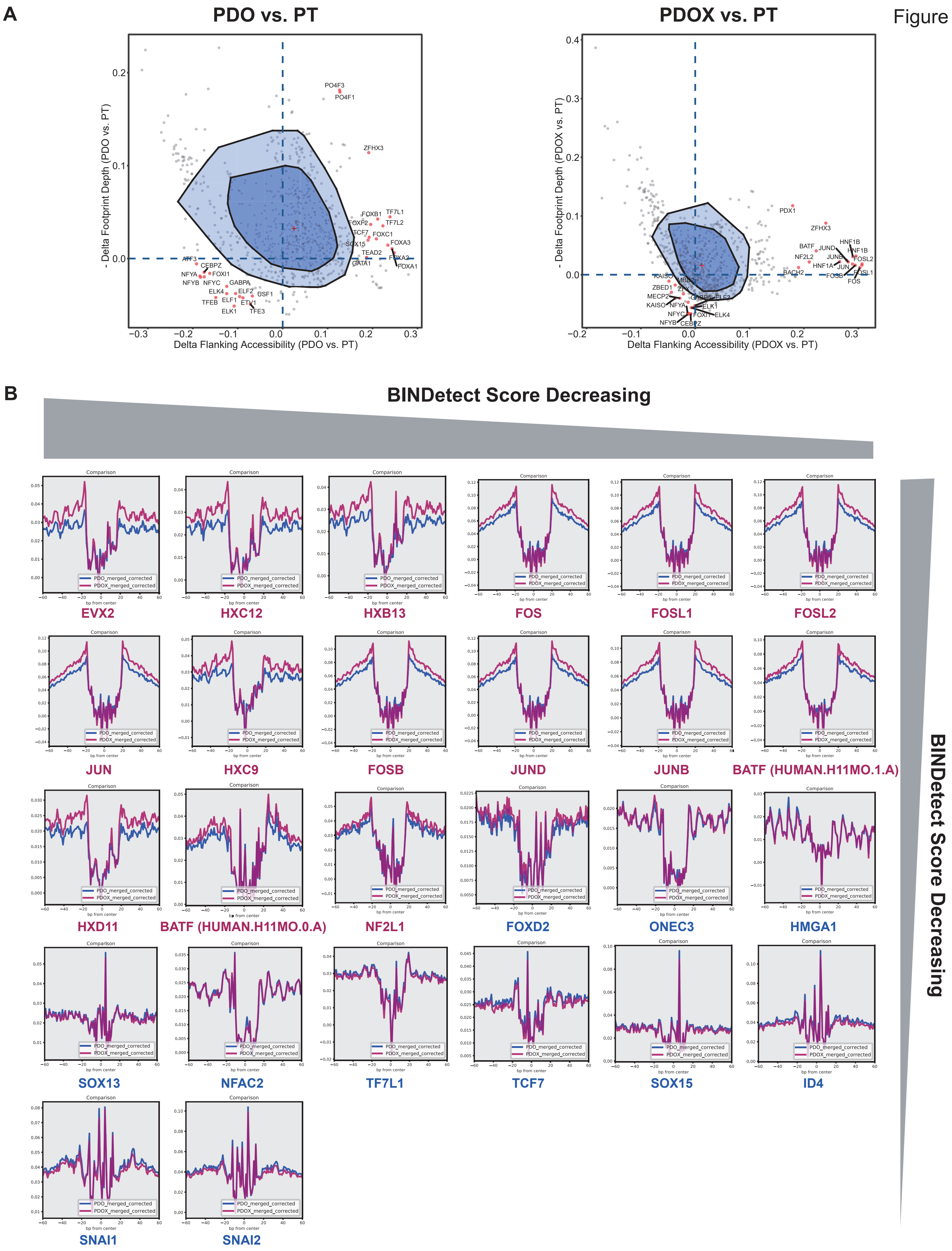
A
PDO vs. PT
PDOX vs. PT
- Delta Footprint Depth (PDO vs. PT)
Delta Flanking Accessibility (PDO vs. PT)
- Delta Footprint Depth (PDOX vs. PT)
Delta Flanking Accessibility (PDOX vs. PT)
B
BINDetect Score Decreasing
BINDetect Score Decreasing
Comparison
PDO_merged_corrected
PDOX_merged_corrected
bp from center
EVX2
HXC12
HXB13
FOS
FOSL1
FOSL2
JUN
HXC9
FOSB
JUND
JUNB
BATF (HUMAN.H11MO.1.A)
HXD11
BATF (HUMAN.H11MO.0.A)
NF2L1
FOXD2
ONEC3
HMGA1
SOX13
NFAC2
TF7L1
TCF7
SOX15
ID4
SNAI1
SNAI2

**Figure S9 Supplemental footprint analysis.**

**(A)** BaGFoot analysis of PDO vs. PT and PDOX vs. PT. The TFs predicted to be active in PDO or PDOX are in the first quadrant beyond the fence area. The TFs predicted to be active in PT are in the third quadrant beyond the fence area. The top 15 TFs in each quadrant based on the distance to the origin are shown.

**(B)** Footprint analysis of the PDOX-active TFs (predicted to be beyond the fence in both comparisons of PDOX vs. PDO and PDOX vs. PT, with the TF names shown in red) and PDO-active TFs (predicted to be beyond the fence in both PDOX vs. PDO and PDO vs. PT, with the TF names shown in blue). For each TF, its footprint in PDOX and PDO are shown. The TFs are ranked by the BINDetect score (EVX2 has the highest absolute value in the active scores in PDOX, and SNAI2 has the highest absolute value in the active scores in PDO).

Figure S10

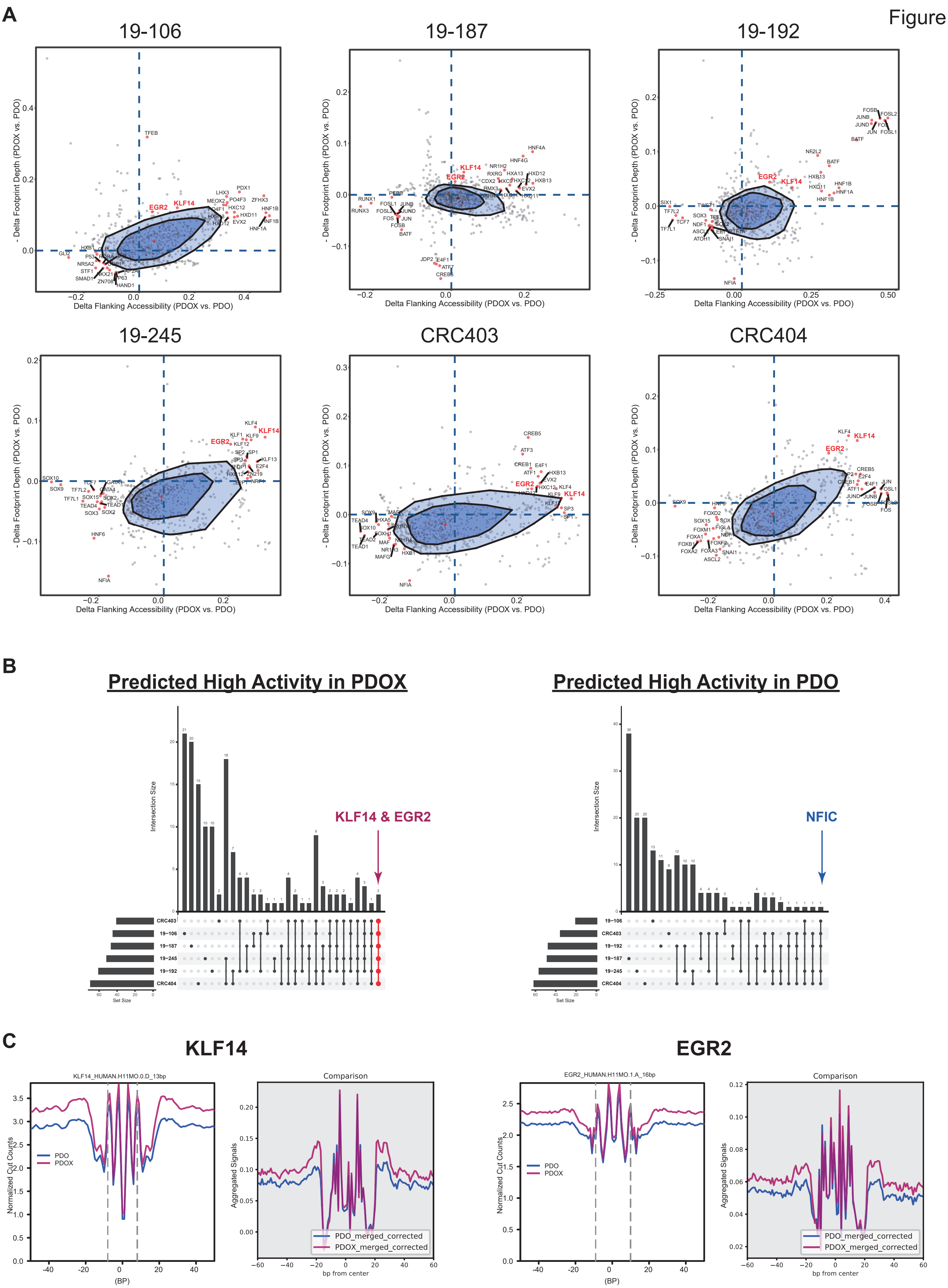
A
19-106
19-187
19-192
19-245
CRC403
CRC404
- Delta Footprint Depth (PDOX vs. PDO)
Delta Flanking Accessibility (PDOX vs. PDO)
KLF14
EGR2
NFIA
B
Predicted High Activity in PDOX
Predicted High Activity in PDO
Intersection Size
Set Size
KLF14 & EGR2
NFIC
CRC403
19-106
19-187
19-245
19-192
CRC404
19-192
C
KLF14
EGR2
KLF14_HUMAN.H11MO.0.D_13bp
EGR2_HUMAN.H11MO.1.A_16bp
Comparison
Normalized Cut Counts
Aggregated Signals
PDO
PDOX
PDO_merged_corrected
PDOX_merged_corrected
(BP)
bp from center

**Figure S10 BaGFoot analysis for individual patient sets.**

**(A)** BaGFoot analysis of PDOX vs. PDO (individual patient set). The top 15 TFs in the first and third quadrants based on the distance to the origin are shown.

**(B)** The upset plots summarizing the overlapped TFs among each individual patient set. KLF14 and EGR2 are predicted to be active in PDOX in all six sets. NFIC is predicted to be active in PDO in five patient sets (all but the 19-187 set).

**(C)** KLF14 and EGR2 footprint comparisons of PDOX and PDO generated by BaGFoot (left for each panel) and BINDetect (right for each panel).

Figure S11

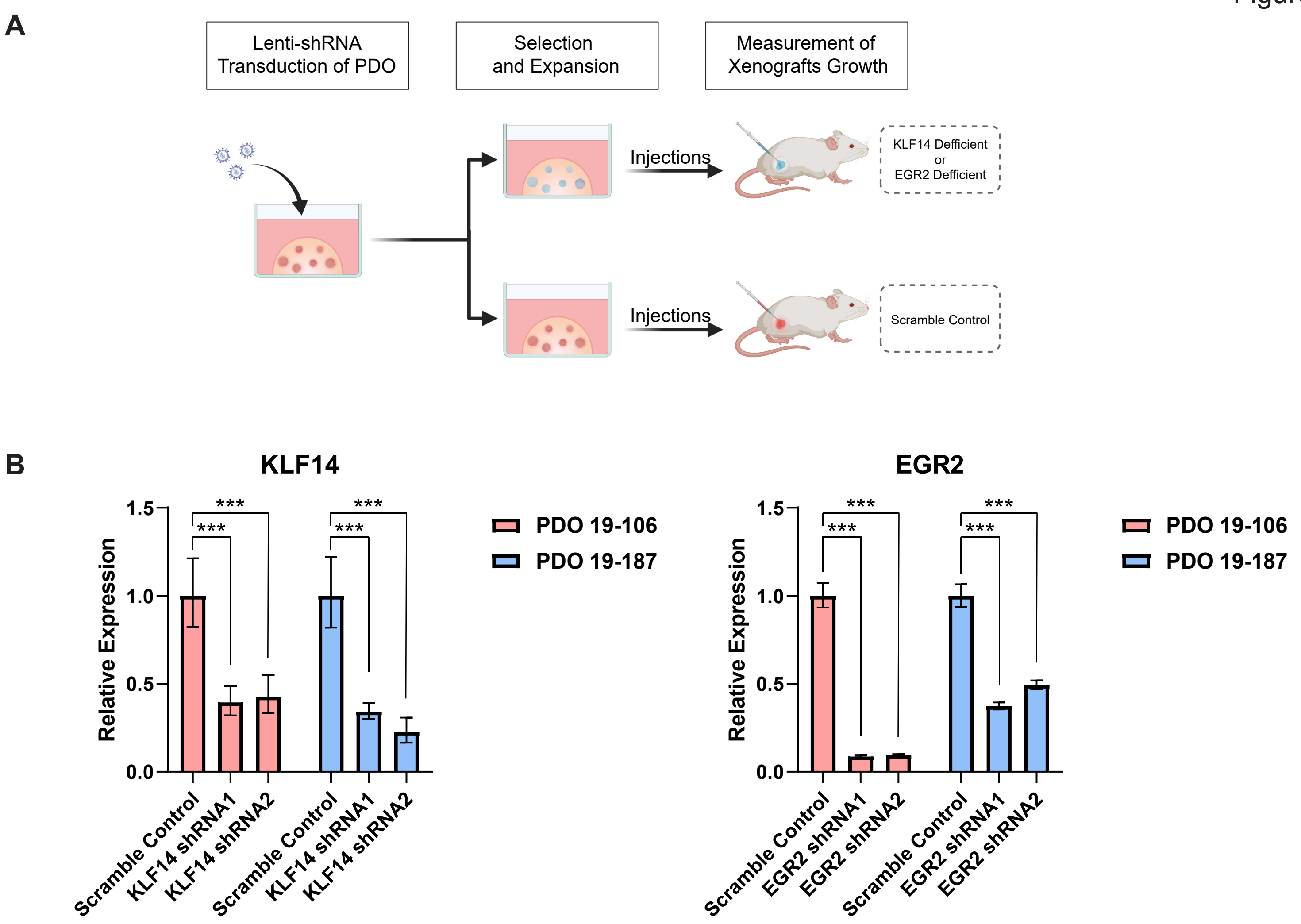
A
Lenti-shRNA Transduction of PDO
Selection and Expansion
Measurement of Xenografts Growth
Injections
KLF14 Defficient or EGR2 Defficient
Injections
Scramble Control
B
KLF14
Relative Expression
0.0
0.5
1.0
1.5
***
***
***
***
Scramble Control
KLF14 shRNA1
KLF14 shRNA2
Scramble Control
KLF14 shRNA1
KLF14 shRNA2
PDO 19-106
PDO 19-187
EGR2
Relative Expression
0.0
0.5
1.0
1.5
***
***
***
***
Scramble Control
EGR2 shRNA1
EGR2 shRNA2
Scramble Control
EGR2 shRNA1
EGR2 shRNA2
PDO 19-106
PDO 19-187

**Figure S11 Supplemental KLF14 and EGR2 *in vivo* validations.**

**(A)** Design of the *in vivo* validation using lenti-shRNA. **(B)** qRT-PCR validated knocking down of KLF14 and EGR2 using the shRNAs. Error bars [relative quantification (RQ) minimum and maximum] indicate 95% confidence interval estimating the mean expressions (N=4). p values were calculated based on ANOVA using $\Delta Ct$, ***$p < 0.001$.

Figure S12

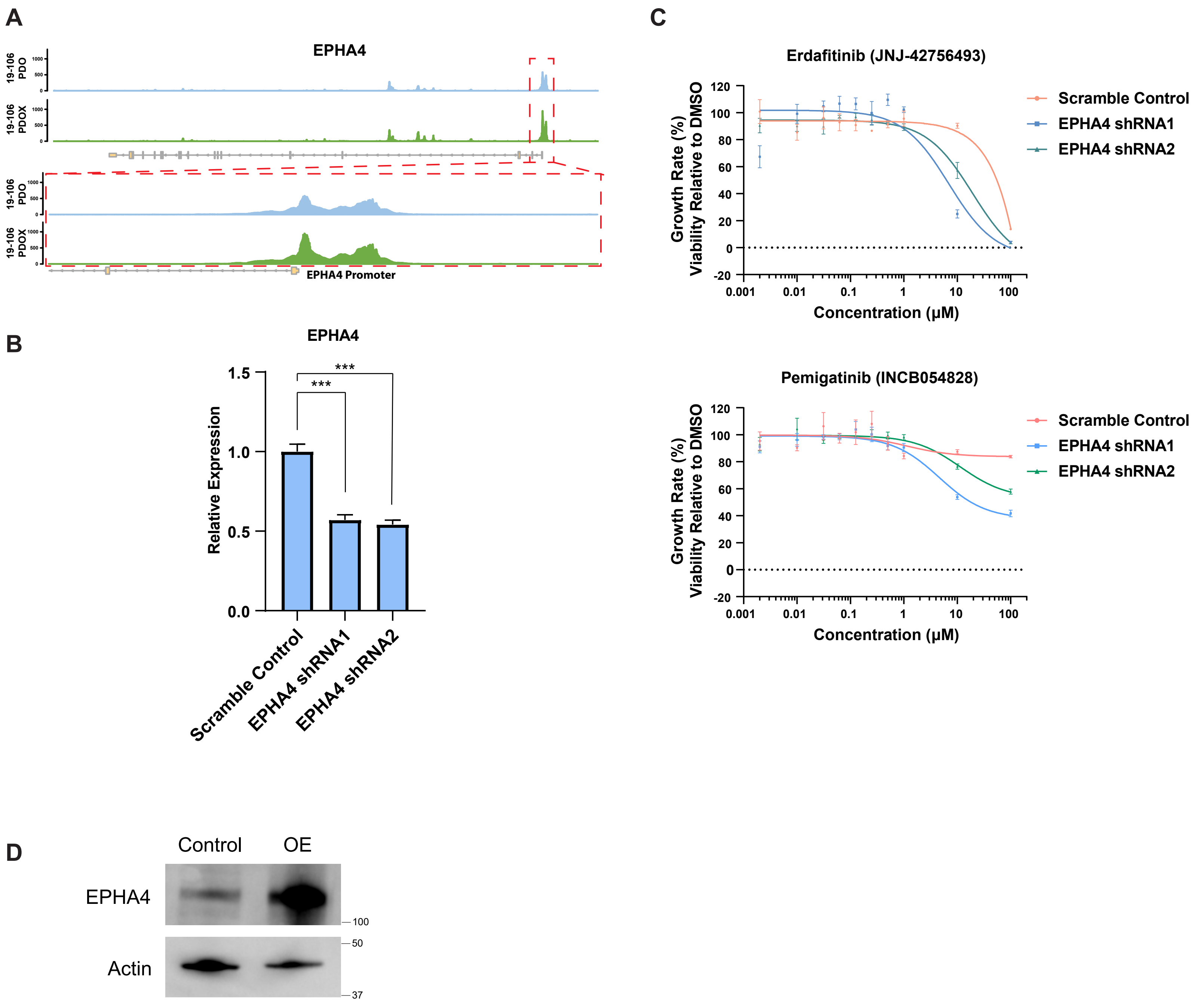
A
EPHA4
19-106 PDO
19-106 PDOX
EPHA4 Promoter
B
EPHA4
Relative Expression
1.5
1.0
0.5
0.0
***
***
Scramble Control
EPHA4 shRNA1
EPHA4 shRNA2
C
Erdafitinib (JNJ-42756493)
Growth Rate (%)
Viability Relative to DMSO
Concentration (μM)
0.001
0.01
0.1
1
10
100
120
100
80
60
40
20
0
-20
Scramble Control
EPHA4 shRNA1
EPHA4 shRNA2
Pemigatinib (INCB054828)
Growth Rate (%)
Viability Relative to DMSO
Concentration (μM)
Scramble Control
EPHA4 shRNA1
EPHA4 shRNA2
D
Control
OE
EPHA4
Actin
100
50
37

**Figure S12 EPHA4 related drug sensitivity tests.**

**(A)** 19-106 ATAC-seq signal track showing EPHA4 locus in PDO and PDOX. The exon locations are indicated in the gene map. The promoter areas of EPHA4 are circled and presented in the bottom panel. **(B)** qRT-PCR validated knocking down of EPHA4 using the shRNAs. Error bars [relative quantification (RQ) minimum and maximum] indicate 95% confidence interval estimating the mean expressions (N=4). p values were calculated based on ANOVA using ΔCt, ***$p < 0.001$. **(C)** PDO19-106 growth rate dose response curves to Erdafitinib and Pemigatinib after knocking down EPHA4. Error bars denote SEM of four replicates. **(D)** Western blot showing EPHA4 overexpression (OE).
